# Whole-blood DNA-methylation patterns during portable-sauna use in firefighters: an exploratory, single-arm pilot study

**DOI:** 10.64898/2026.09.02.748979

**Authors:** Varun B. Dwaraka, Adiv A. Johnson, Kirsten Seale, Nolan Kahal, Dua Sheikh, Sean Morrissey, Ryan Smith

## Abstract

Research in model organisms has consistently shown that mild, non-lethal heat stress can prolong lifespan. Although causal data are lacking in more complex animals, human observational research has linked sauna bathing to cardiovascular protection and improved age-related outcomes. To examine whether sauna use may influence aging-related biology, we retrospectively analyzed longitudinal whole-blood methylomic data from 18 firefighters who used a portable sauna. A separately processed cohort of 36 matched adults provided descriptive context; firefighters showed higher DNA methylation– estimated maximal oxygen uptake (DNAmVO2 max), lower AdaptAge, and higher DamAge. After 8–18 weeks, eight predictors met a within-family Benjamini–Hochberg false-discovery-rate (FDR) threshold of <0.05 underage/sex-adjusted epigenetic age acceleration: PCDNAmTL, DepressionBarbu, SystemsAge Blood, SystemsAge Lung, DNAmPulsePressure, DNAmHDL, DNAmCystatinC, and DNAmDHEAS. DNAmDHEAS increased, whereas the other predictors changed in directions consistent with short-term findings for high-intensity interval training, another hormetic stressor. None met the within-family FDR threshold after adjustment for 12-cell leukocyte composition, although effect estimates were not uniformly eliminated. At individual cytosine-phosphate-guanine (CpG) sites, BACON empirical-null calibration retained 126 acute and 329 long-term candidates at P<0.0001; none was FDR-significant. DMRcate identified five acute and 140 long-term differentially methylated regions using a harmonic mean of individual CpG FDR values <0.05. Long-term gometh analysis also identified ATP-binding cassette (ABC) transporter enrichment (FDR=0.026). These exploratory findings suggest that portable-sauna use may influence the blood methylome within 8–18 weeks while underscoring the importance of leukocyte-composition and technical sensitivity analyses.

## Introduction

An intriguing relationship between heat stress and longevity exists in simpler model organisms. In the nematode *Caenorhabditis remanei*, Chen and Maklakov showed that repeated heat-induced mortality led to the evolution of longer lifespans [1]. Conversely and in line with the evolutionary theories of aging [2], random extrinsic mortality selected for the evolution of shorter lifespans [1]. Non-lethal stress has been shown to prolong lifespan in *Drosophila melanogaster* [3] and elevate the expression of the heat-shock protein Hsp70 [4]. *Caenorhabditis elegans* worms overexpressing the naked mole-rat version of *Hspb1* (encoding for Heat shock protein beta-1) are longer-lived and more resistant to heat. Moreover, their prolonged lifespan requires the activity of *hsf-1* (encoding for Heat shock transcription factor hsf-1) [5].The overexpression of *hsf-1* itself boosts longevity in worms and does so via the renovation of mitochondrial networks [6]. Heat-induced life extension has also been documented in yeast [7].

While causal evidence in more complex animals is lacking, there is a surfeit of correlational evidence in humans. Specifically, a large body of literature has associated sauna usage with improved health outcomes [8]. In one prospective cohort study, Laukkanen et al followed 2,315 middle-aged Finnish men for close to 21 years. For those that engaged in sauna bathing two to three times each week, the hazard ratio for sudden cardiac death was 0.78. The hazard ratio dropped down to 0.37 for the group engaging in sauna bathing four to seven times each week. Beneficial associations were also observed for all-cause mortality, mortality due to cardiovascular disease, and fatal coronary heart disease. Session duration was also relevant, as outcomes improved as the length of time increased from under 11 minutes to more than 19 minutes [8, 9]. Mechanistically, sauna bathing has been hypothesized to enhance cardiovascular health by decreasing arterial stiffness, improving lipid biomarkers, reducing blood pressure, enhancing endothelium-mediated dilation, and modulating the autonomic nervous system [8].

To better understand the relationship between sauna bathing and aging in humans, we retrospectively analyzed whole blood Illumina Infinium MethylationEPIC v2.0 data from 18 firefighters that participated in a exploratory pilot study involving a portable sauna. Samples were obtained at baseline, approximately one hour after one session, and after 8-18 weeks. In addition to analyzing a variety of epigenetic aging clocks and epigenetic biomarker proxies, we looked at CpG-level changes across timepoints. Finally, these 18 firefighters were compared to a matched, contextual cohort. This work, which represents the first study investigating the effects of sauna bathing on epigenetic clocks and proxies, provides initial evidence of epigenetic remodeling during portable-sauna use and identifies candidate loci for further investigation.

## Materials and Methods

### Participants and design

This manuscript reports a retrospective, exploratory analysis of prospectively collected samples and data from a single-arm pilot portable-sauna program. Nineteen firefighters enrolled in an eight-week home program (approximately 20 minutes per session, with an instructed frequency of at least four sessions per week). Eighteen participants had analyzable EPICv2custom whole-blood methylation (15 males and three females; mean age = 42.6 years; age range = 24.9-54.8 years). No a priori power calculation or single primary methylation endpoint was specified for this secondary analysis. Samples were collected at timepoint 1 (TP1; pre-sauna baseline), timepoint 2 (TP2 or acute; approximately one hour after a single session and same day), and timepoint 3 (TP3 or long-term; after 8-18 weeks of regular use). Two paired within-subject contrasts were defined for this exploratory analysis: acute (TP1 vs TP2, n=10 pairs) and long-term (TP1 vs TP3, n=11 pairs).

Because the baseline and post-session (TP1/TP2) samples were collected on the same calendar day for most participants, within-subject pre/post order was resolved from LIMS sample-collection timestamps (time of day) rather than sample-processing/plating order. The timestamps showed that four participants’ second draws were not collected within the same sauna session (27.5 hours to 59 days after baseline). These draws were excluded from the acute contrast; three of the four participants had a valid long-term (TP3) sample and were retained in the long-term contrast, while the fourth (whose second draw was 59 days post-baseline) was excluded entirely, consistent with the criteria used across both this pipeline and an independent collaborator re-analysis.

Device setting and adherence/tolerability were captured through hydration and symptom logs (Figure S5; Tables S15-S16). A total of 188 form entries were submitted. Individually linkable adherence counts were available for eight of the 11 long-term participants and summed to 121 sessions. Chamber temperature, relative humidity, skin temperature, and core temperature were not continuously measured; the recorded 0-7 device setting is therefore not treated as a calibrated physiological heat dose. Quality of life was assessed with the WHO-QoL instrument at baseline (Figure S4; Table S14). The reference cohort comprised 36 disease-free, age/sex-matched adults from the TruDiagnostic normative survey database (12-cell EPICv2Custom). Participants who reported no health conditions were matched without replacement to firefighters by sex and an age caliper of 3 years in either direction (k=2). Cohort characteristics and design are summarized in Table 1 and Figure 1.

**Fig. 1.**
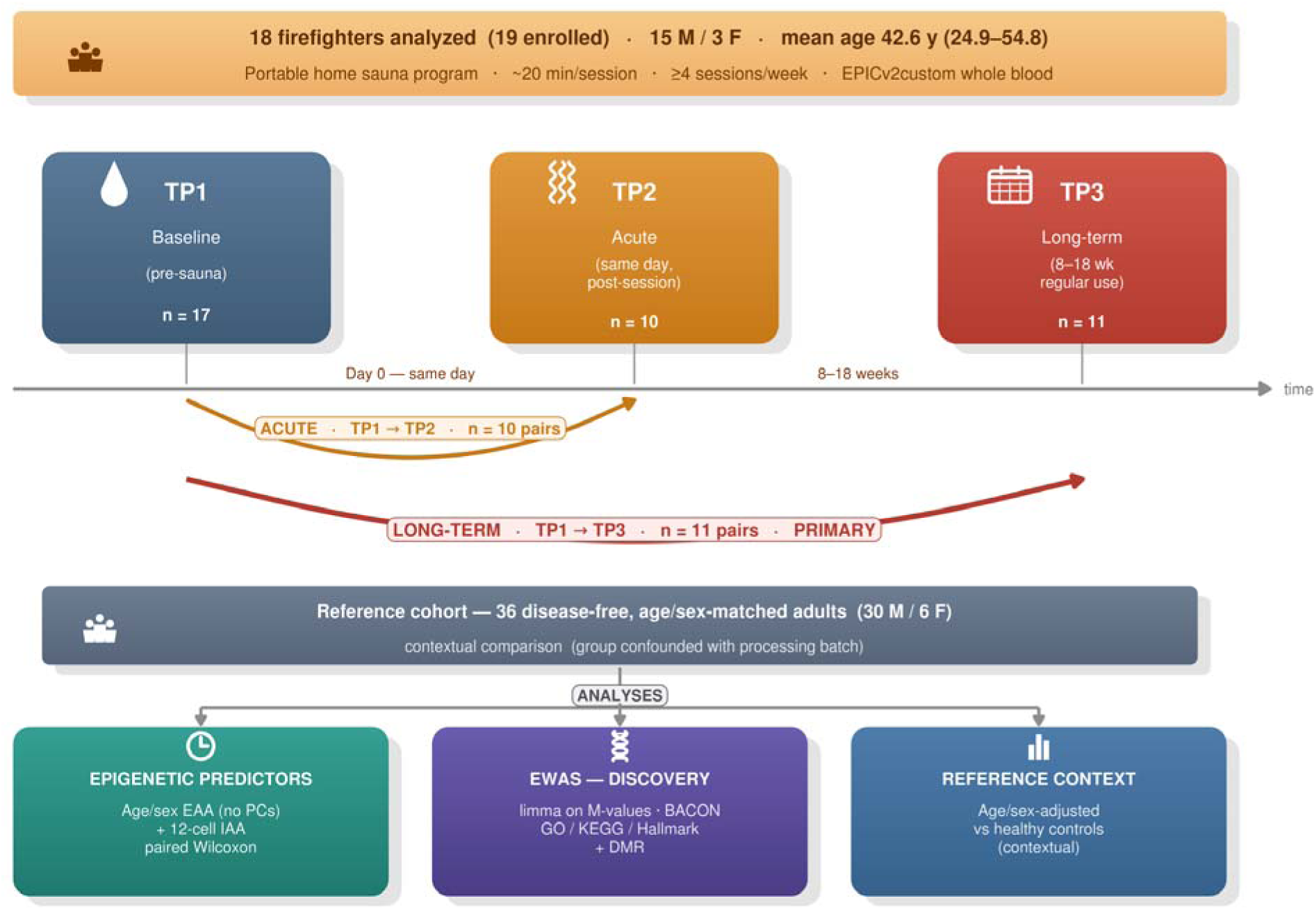
Study design and cohort overview. Nineteen firefighters enrolled and 18 had analyzable EPICv2custom whole-blood DNA methylation (15 men and 3 women; mean age = 42.6 years; age range = 24.9 -54.8 years). Samples were available at baseline (TP1, n=17), after a single same-day portable-sauna session (TP2, n=10), and after 8–18 weeks of regular home use (TP3, n=11). Paired analyses comprised 10 acute TP1–TP2 pairs and 11 long-term TP1–TP3 pairs. The instructed program was approximately 20 minutes per session at least four times weekly. A separately processed cohort of 36 disease-free, age/sex-matched adults was used for contextual comparison. The lower panels summarize the predictor, epigenome-wide/pathway/DMR, and reference-cohort analyses

**Table 1.** Cohort demographics and study characteristics. Enrollment and analyzable sample counts, sex and age distributions, samples and paired contrasts by timepoint, participant-linked adherence, recorded heat levels, submitted hydration/symptom entries, and characteristics of the separately processed reference cohort.

| Characteristic | Value |
| --- | --- |
| Firefighters enrolled / analyzed | 19 / 18 |
| Sex (M / F) | 15 / 3 |
| Age, mean (range), y | 42.6 (24.9-54.8) |
| Samples at TP1 (baseline) | 17 |
| Samples at TP2 (acute) | 10 |
| Samples at TP3 (long-term) | 11 |
| Acute paired contrast (TP1->TP2), n pairs | 10 |
| Long-term paired contrast (TP1->TP3), n pairs | 11 |
| Long-term paired contrast sex (M / F) | 10 / 1 |
| Sessions logged per participant, median (range) | 15.5 (1-32) |
| Participants with individually linkable adherence records | 8 of 11 long-term participants |
| Individually linked sessions, total | 121 |
| Sauna heat level (0-7), median (IQR) | 6 (5-6) |
| Submitted hydration/symptom form entries | 188 |
| Symptom reports among submitted entries | 2 (1 headache, 1 dry mouth) |
| Reference cohort (disease-free, matched) | 36 |
| Reference sex (M / F) | 30 / 6 |
| Reference age, mean (range), y | 42.8 (24.9-54.8) |
| Reference age match (Wilcoxon p) | 0.847 |

### Methylation processing

IDAT files were processed with the TruDiagnostic IRC pipeline. EPICv2custom data were normalized by single-sample Noob (ssNoob; minfi) [10] with detection-p quality control, and missing values for clock CpGs were imputed with SeSAMe [11]. Because ssNoob is a single-sample method, firefighter and control predictors were computed in separate runs (equivalent to co-normalization and required to avoid a 79-sample memory limit). Beta values were used for reporting, and M-values (logit2) were used for EWAS. Per-sample QC (normalized beta-value distributions, central tendency) is shown in Figure S1.

### Epigenetic-age acceleration (EAA/IAA)

For every predictor, EAA = residuals(predictor ∼ Age + Sex) and IAA = residuals(predictor ∼ Age + Sex + 11 raw immune-cell-proportion covariates), both fit within the 18 firefighters only. Control-probe principal components were excluded from both: in this longitudinal design PC1 was confounded with timepoint (all 11 long-term subjects shifted PC1 in the same direction; Figure S2). Adjustment for this axis could remove both technical and intervention-associated variation, whereas exclusion could retain both. The no-PC predictor models therefore describe timepoint-associated change and are not assumed to uniquely identify an intervention effect. Within the 11 paired long-term participants, TP1 and TP3 samples occupied disjoint sets of physical Illumina BeadChips, and 0 of 11 participants had baseline and follow-up samples on the same chip (Table S17; Figure S8). Thus physical-chip assignment and long-term timepoint were completely confounded in the paired contrast; no model fitted to these array data can estimate a long-term timepoint coefficient independently of chip.

Two further design choices in the IAA model merit explicit justification, since IAA is the primary readout for the acute contrast. First, the age/sex-matched reference cohort, which was used elsewhere for contextual comparison (Table S9), was excluded from the IAA fitting sample. The reference cohort and firefighters were normalized in entirely separate ssNoob runs (see *Methylation processing*), so cohort membership is completely confounded with processing batch for that comparison; no regression can attribute a covariate effect to biology rather than batch when the two never vary independently. Permutation-based methods exist for batch correction when batch is confounded with a grouping of interest (e.g., permuted surrogate variable analysis) [12], but they are designed for larger cross-sectional cohorts and are unnecessary here: the reference cohort is not part of the primary within-subject contrast, so the confound is avoided by simply restricting IAA fitting to the 18 firefighters, among whom no such batch/cohort confound exists.

Second, immune adjustment used the 12-cell deconvolution (EpiDISH/Teschendorff; CD4Tnv/mem, CD8Tnv/mem, Baso, Bmem, Bnv, Treg, Eos, NK, Neu, Mono) [13] entered as 11 raw proportions (Monocytes as implicit reference). These proportions are compositionally constrained (sum to approximately 1) and produce an ill-conditioned design matrix (condition number approximately 4.3 x 10^4^); however, the matrix remains full rank with no aliased coefficients, and collinearity inflates individual coefficient standard errors without biasing the fitted values or residuals from which IAA was computed. Residuals from the raw 11-covariate model were numerically identical, to floating-point precision, to a full/untruncated 11-component PCA rotation of the same covariates. A reduced-dimension (PCA-truncated) version of this adjustment was evaluated and rejected: truncating to a smaller number of principal components is a real, discretionary modeling choice, and the resulting IAA hit count proved highly unstable across plausible choices (ranging from 0 to >100 significant predictors depending on how many components were retained, with no natural stopping point and no improvement in stability under standard objective component-selection rules). The raw, untruncated covariate set was retained instead, since it requires no such choice. Sensitivity of the cell-composition-adjusted results to deconvolution reference was separately assessed with a 19-cell CAB-refined panel (also entered raw, unreduced; Figure S6; Table S12): largely consistent with the 12-cell model, with one exception: DNAmCystatinC reaches IAA-FDR<0.05 under the 19-cell reference (FDR=0.0098) but not the primary 12-cell reference (FDR=0.52). All 11 subjects shifted in the same direction under that sensitivity model, but the result is reported as sensitivity-dependent rather than a primary finding. Several predictors (pcgtAge, irS, irS2, tnsc2/tnsc, AdaptAge) remain nominally significant under both references (p=0.007-0.032, not FDR-significant either way). These results were directionally consistent with, but not identical to, the 12-cell result, and were reported as a sensitivity check rather than a discrepancy to be resolved.

### Predictor panels

The predictor panels included PC clocks (PCHorvath1/2, PCHannum, PCPhenoAge, PCGrimAge, and PCDNAmTL); DunedinPACE, PhenoAge, and OMICmAge; SystemsAge organ clocks; DNAm PhysAge physiological proxies (DNAmDHEAS, DNAmHDL, DNAmHbA1c, DNAmCystatinC, DNAmPulsePressure, and others); intrinsic-rate (irS/irS2) and mitotic (tnsc2/tnsc) indices; and additional standalone clock and behavioral-exposure methylation scores, including AdaptAge, DamAge, CausAge, ENCen100, ENCen40, DepressionBarbu, IntrinClock, and PedBE. A prior pass omitted several predictors because the seed list was drawn from a legacy pipeline report rather than the full set of predictors computed by the pipeline. We corrected this issue by scanning the raw predictor set directly and applying content-based exclusions established independently of the results. Excluded variables included environmental-exposure scores, technical PCs, duplicate encodings, decomposition sub-scores of an already tested composite, and one clock superseded by a newer version already in the panel. The corrected panels comprised 83 candidate Clocks-family predictors, of which 81 yielded analyzable paired tests and entered the within-family FDR correction (up from 35 tested previously). The irT and irT2 predictors were constant population-median scalars and therefore could not produce paired tests. The panels also included 113 Marioni protein EpiScores (up from 69) and EBP v1 and v2 predictors. The EBP v2 metabolite-proxy panel was recomputed with the same age/sex EAA and 12-cell IAA procedures used for the other predictors. Predictor results are reported in Tables 2-4 (significant, tiered) with the full unfiltered panels in Tables S1-S4 (clocks, PhysAge, EBP v2, Marioni); predictor response is visualized in Figure 2 and the reference comparison in Figure 3.

**Fig. 2.**
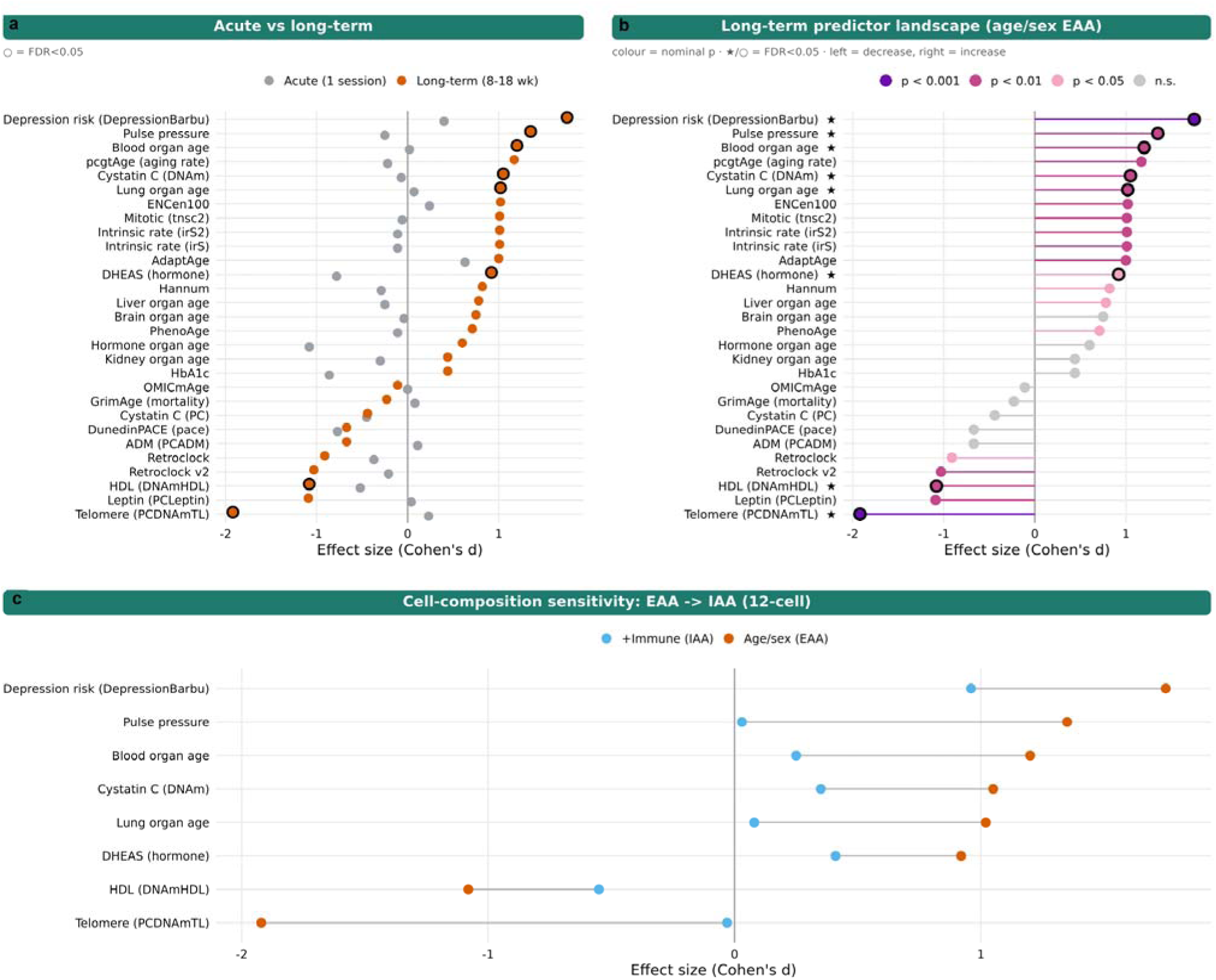
Epigenetic predictor response. (a) Selected acute and long-term standardized paired effect sizes (Cohen’s d) for epigenetic clocks and methylation-derived predictors; black-outlined points met within-family Benjamini–Hochberg FDR<0.05. (b) Selected long-term age/sex-adjusted epigenetic age acceleration (EAA) results, colored by nominal-p tier; stars and outlined points identify within-family FDR<0.05. (c) Sensitivity of the eight long-term EAA findings that met within-family FDR<0.05 to adjustment for the primary 12-cell methylation-inferred leukocyte reference (IAA). None of the eight remained at within-family FDR<0.05 after 12-cell adjustment, although effect estimates were not uniformly eliminated. Positive and negative values indicate the direction of the methylation-derived score and are not a uniform favorable-versus-unfavorable axis. Predictor EAA and IAA models did not include control-probe PCs. Source statistics are in Tables 2–3 and Supplementary Tables S1–S2

**Fig. 3.**
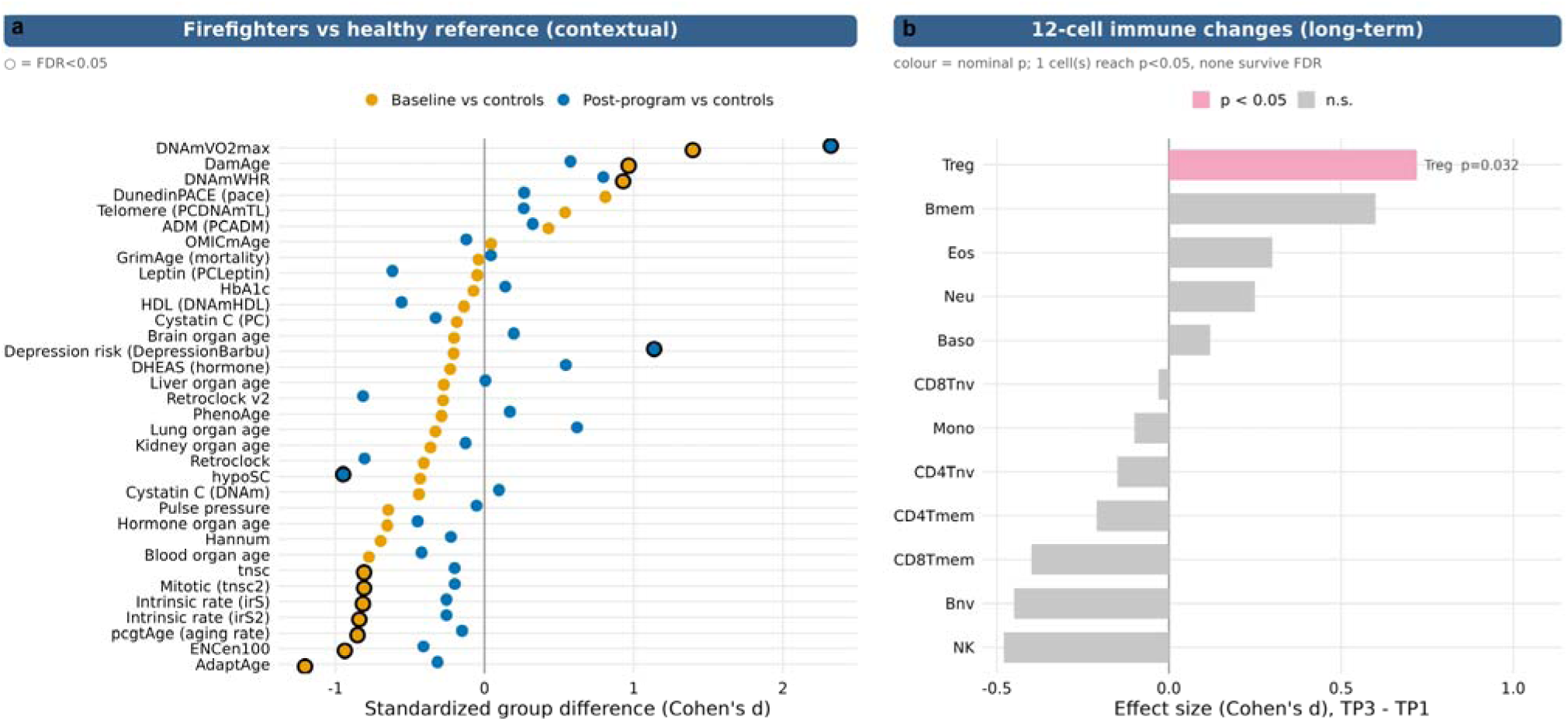
Reference-cohort context and methylation-inferred leukocyte changes. (a) Selected standardized differences (Cohen’s d) between firefighters and the separately processed disease-free reference cohort at baseline and long-term follow-up; outlined points met within-family BH-FDR<0.05. These comparisons are descriptive because cohort and processing run are aligned. (b) Standardized paired TP3-minus-TP1 changes for 12 methylation-inferred leukocyte proportions among 11 long-term pairs. Regulatory T cells provided the strongest individual signal (d=0.72, p=0.0322, BH-FDR=0.386); no cell proportion met BH-FDR<0.05. Source statistics are in Supplementary Tables S9 and S11

**Table 2.**
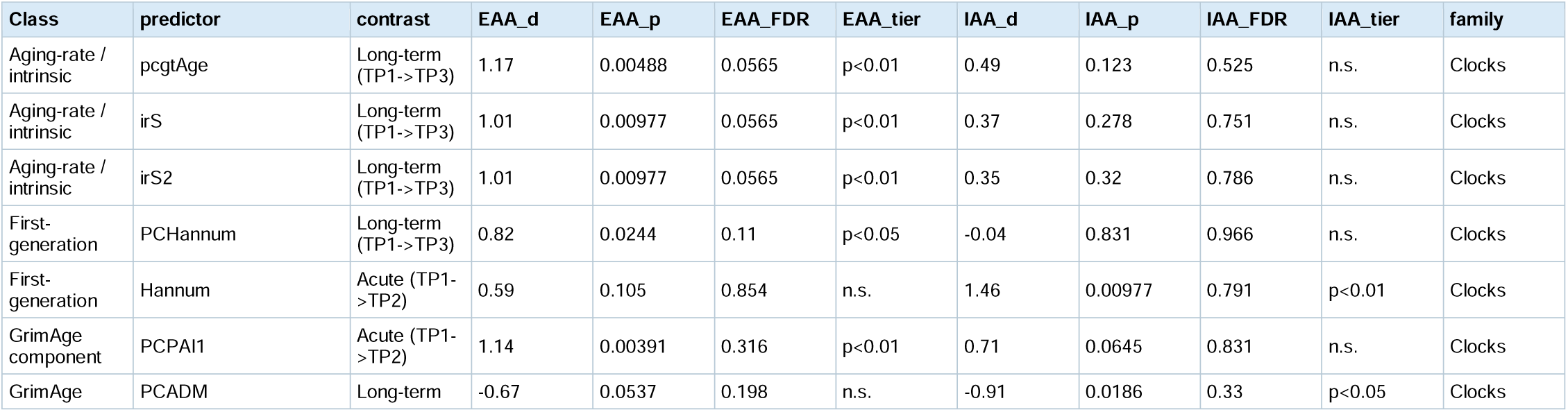
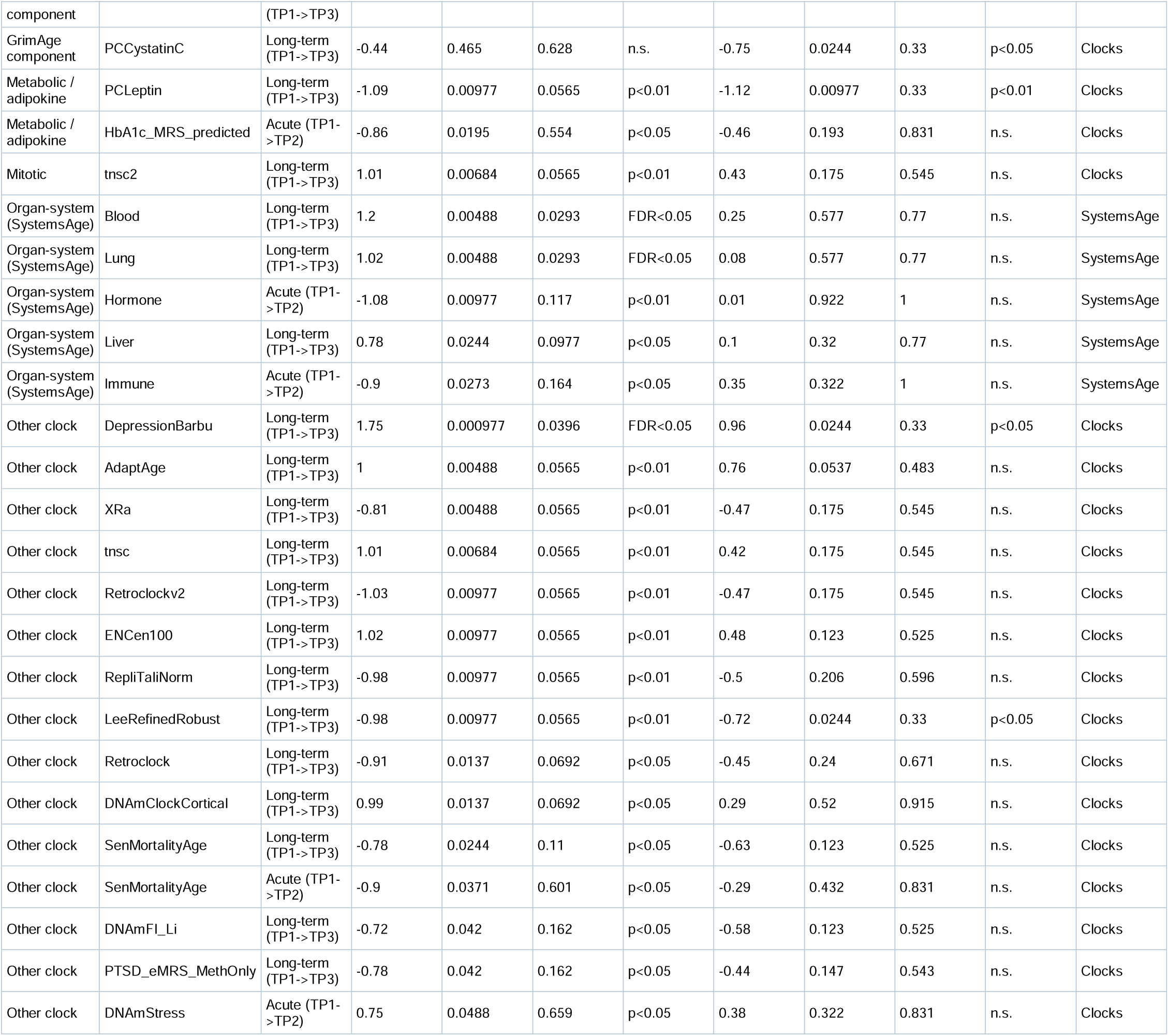
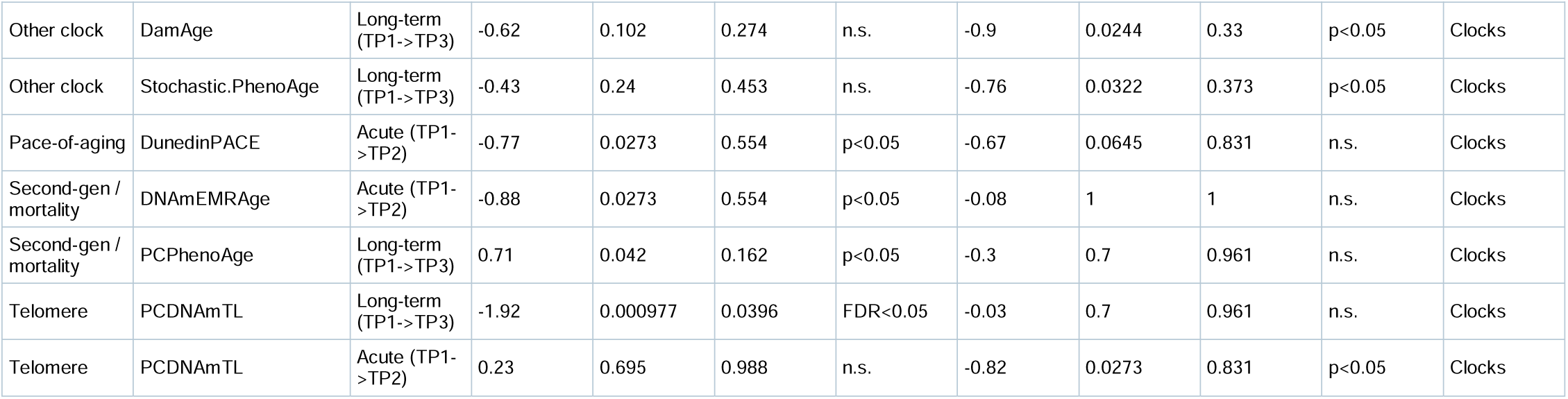
Selected epigenetic clocks and SystemsAge organ predictors. The table contains the acute or long-term results with nominal p<0.05 under age/sex-adjusted EAA or primary 12-cell IAA (acute and long-term), with Cohen’s d, nominal p values, within-family BH-FDR values, and significance tiers. The complete predictor panel for both contrasts is in Supplementary Table S1.

**Table 3.** Selected DNAm PhysAge subcomponents. The table contains the acute or long-term physiological-trait methylation proxies with nominal p<0.05 under age/sex-adjusted EAA or primary 12-cell IAA (one acute and five long-term), with Cohen’s d, nominal p values, and within-family BH-FDR values. These are methylation-derived proxies rather than direct clinical measurements. The complete 10-predictor panel for both contrasts is in Supplementary Table S2.

| family | predictor | EAA_d | EAA_p | IAA_d | IAA_p | EAA_FDR | IAA_FDR | contrast | EAA_tier | IAA_tier |
| --- | --- | --- | --- | --- | --- | --- | --- | --- | --- | --- |
| PhysAge | DNAmPulsePr_final | 1.35 | 0.00488 | 0.03 | 0.765 | 0.0228 | 1 | Long-term (TP1->TP3) | FDR<0.05 | n.s. |
| PhysAge | DNAmHDL_final | -1.08 | 0.00488 | -0.55 | 0.102 | 0.0228 | 0.515 | Long-term (TP1->TP3) | FDR<0.05 | n.s. |
| PhysAge | DNAmCystatinC_final | 1.05 | 0.00684 | 0.35 | 0.206 | 0.0228 | 0.515 | Long-term (TP1->TP3) | FDR<0.05 | n.s. |
| PhysAge | DNAmDHEAS_final | 0.92 | 0.0137 | 0.41 | 0.206 | 0.0342 | 0.515 | Long-term (TP1->TP3) | FDR<0.05 | n.s. |
| PhysAge | DNAmDHEAS_final | -0.78 | 0.0195 | -0.41 | 0.322 | 0.195 | 1 | Acute (TP1->TP2) | p<0.05 | n.s. |
| PhysAge | DNAmCRP_final | 0.78 | 0.042 | 0.49 | 0.206 | 0.084 | 0.515 | Long-term (TP1->TP3) | p<0.05 | n.s. |

**Table 4.**
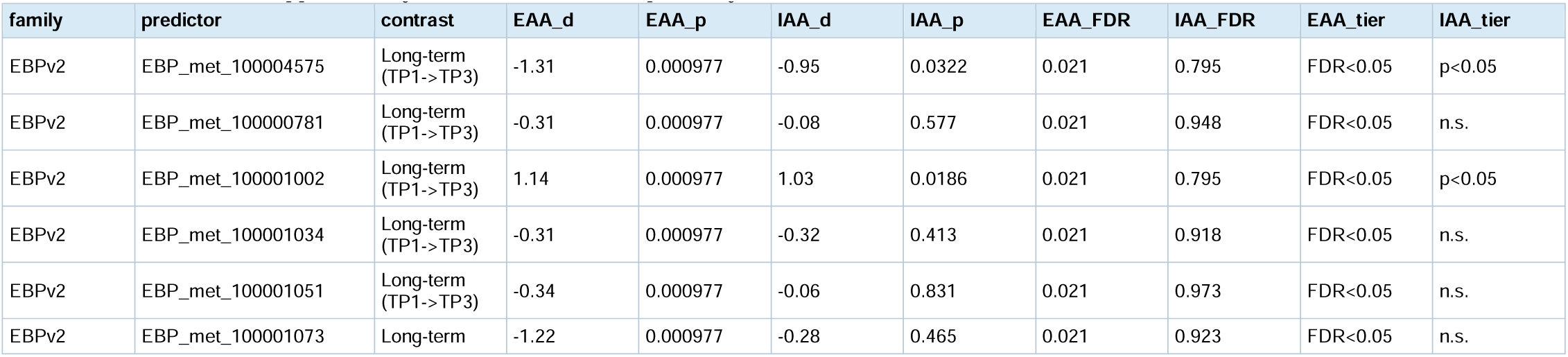
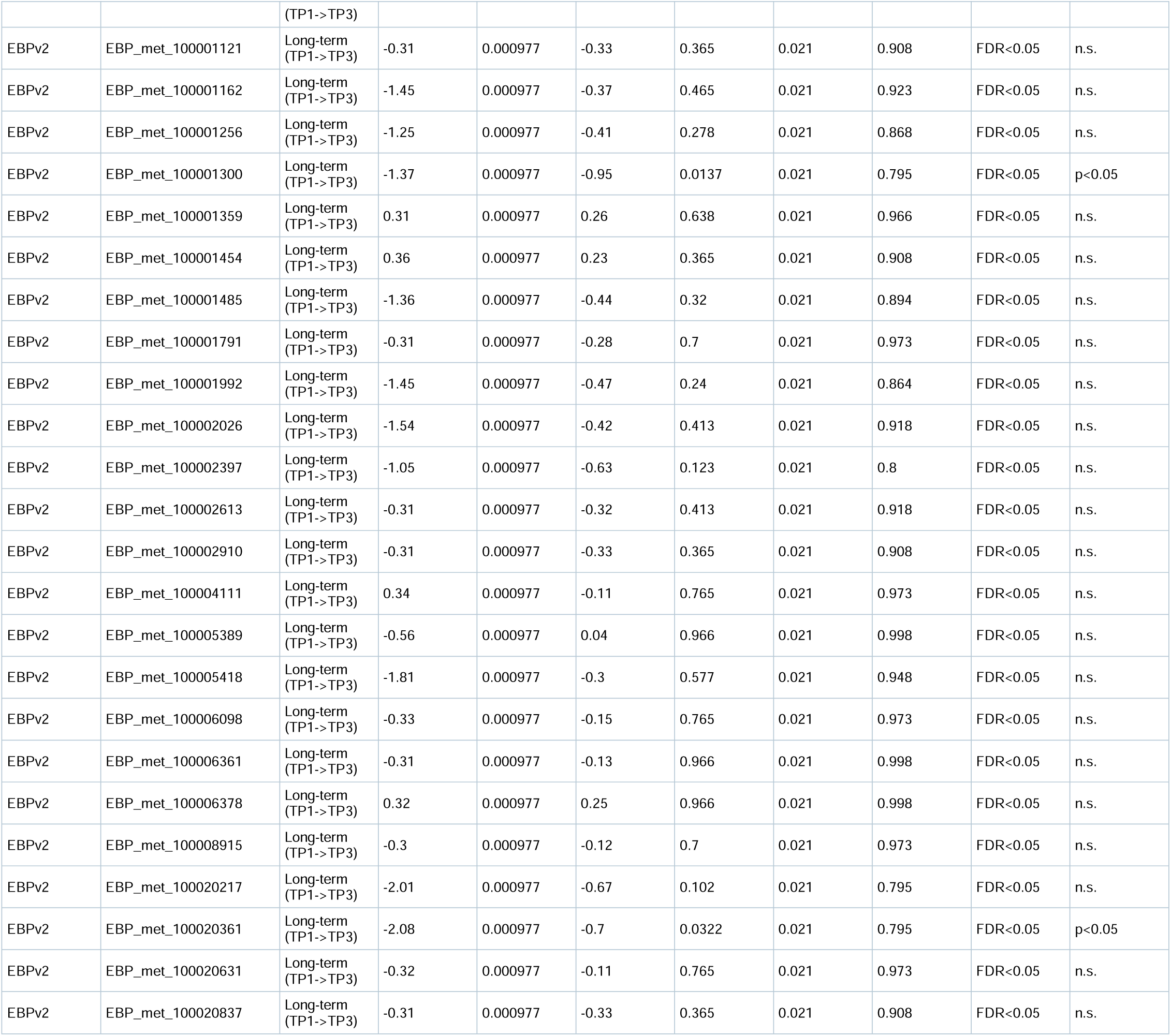
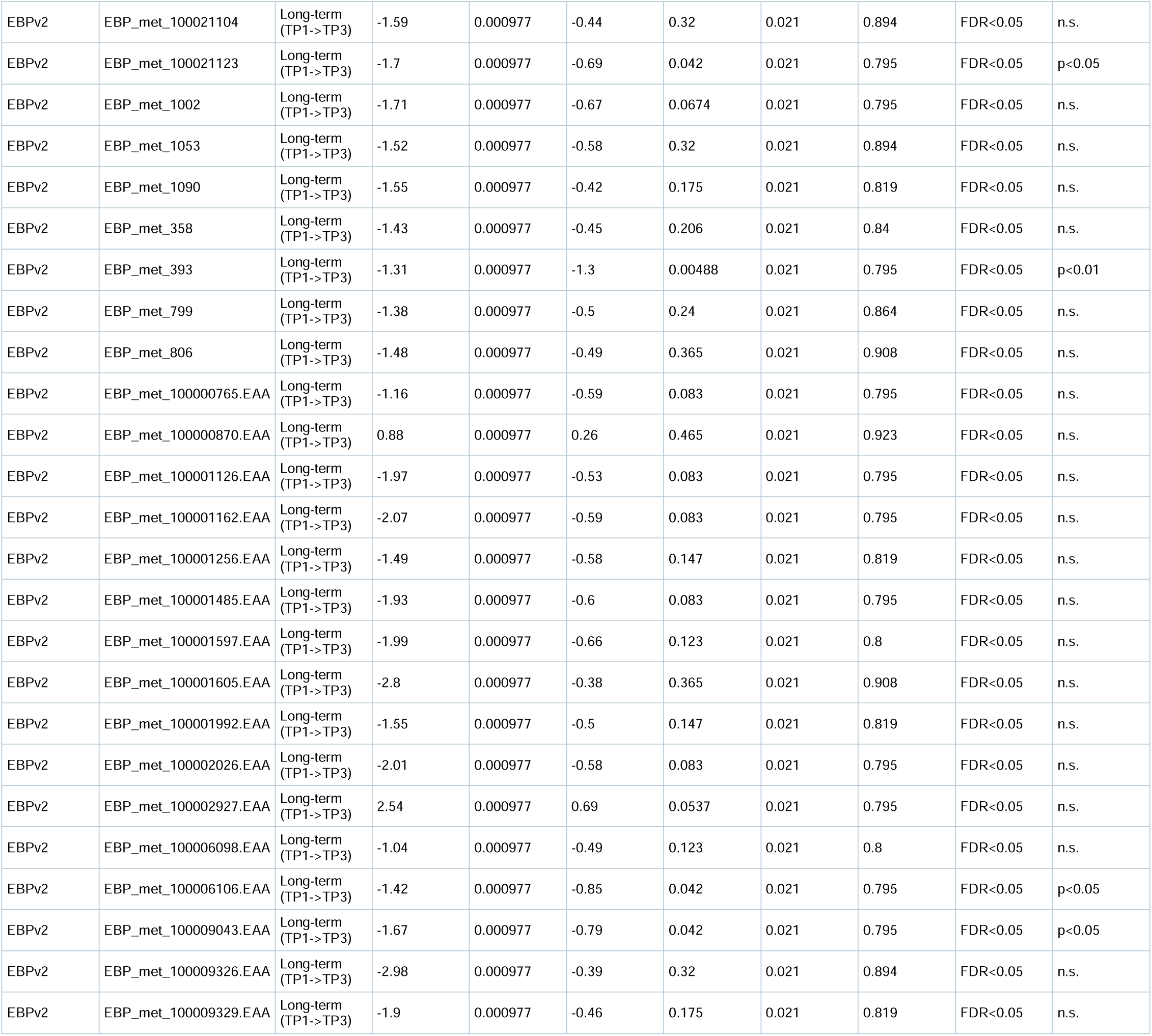
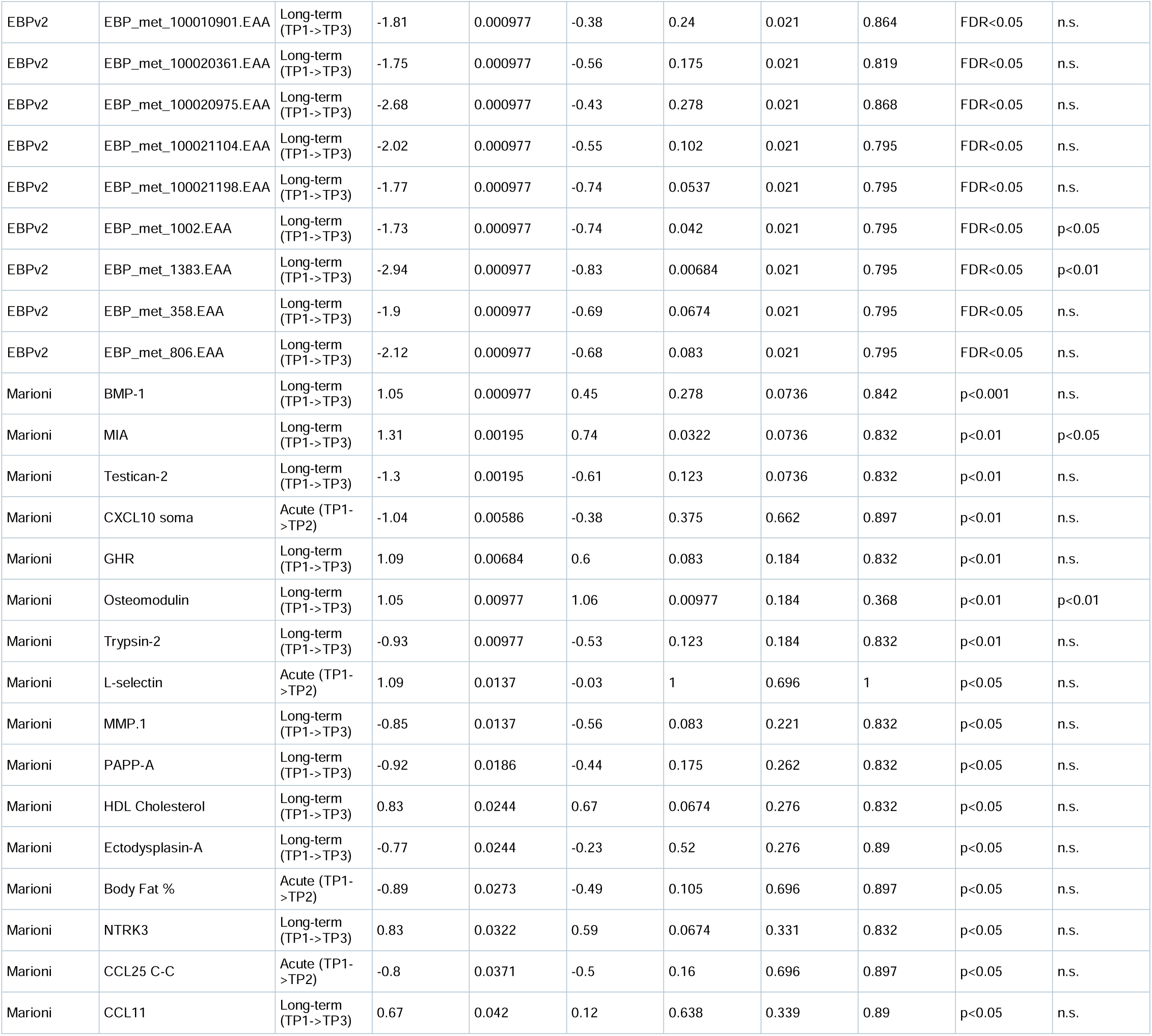
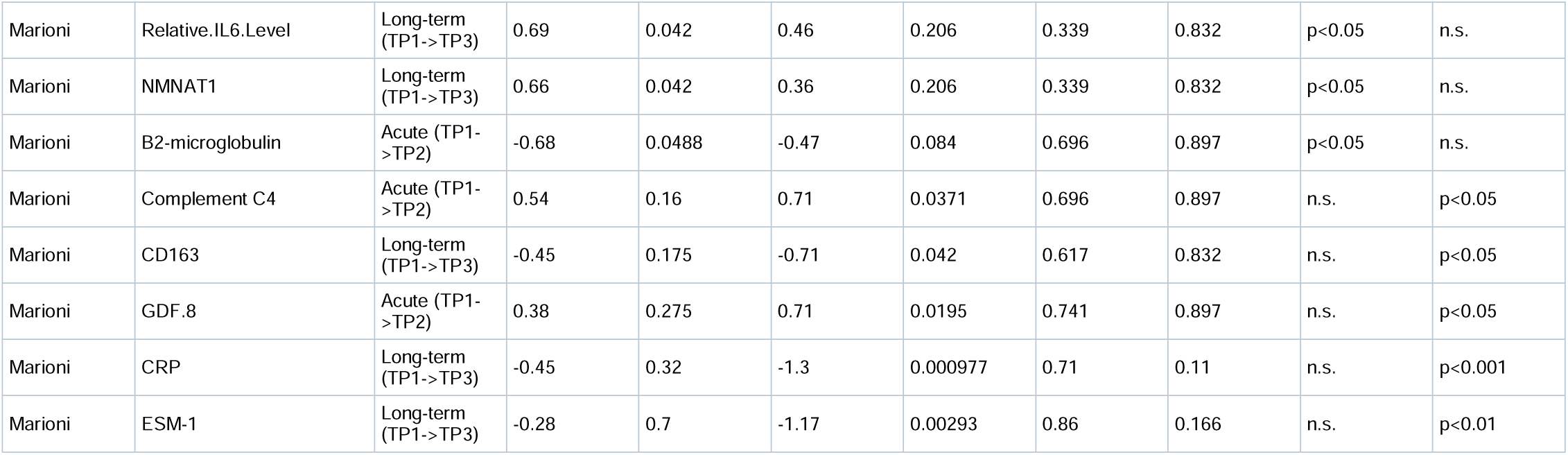
Selected EBP v2 metabolite-proxy and Marioni protein EpiScore results. The table contains long-term EBP v2 scores at EAA p=0.000977 (within-family BH-FDR=0.021) and 24 Marioni EpiScores with nominal p<0.05 under EAA or IAA (7 acute and 17 long-term), for 88 displayed rows. No Marioni EpiScore met within-family BH-FDR<0.05. Complete EBP v2 and Marioni results for both contrasts are in Supplementary Tables S3 and S4, respectively.

### Statistics: predictors

Paired within-subject contrasts were tested by the exact Wilcoxon signed-rank test, with Cohen’s d used as the effect-size measure and Benjamini-Hochberg FDR controlled within each predictor family. Given n=10-11 (exact p-floor approximately 0.001), within-family BH-FDR is achievable for the focused clock/organ/functional panels but not for the large Marioni/EBP families; those are reported using nominal p values and effect sizes (exploratory). Significant results are tabulated with a tier column (strongest of unadjusted p<0.05 / p<0.01 / p<0.001 / FDR<0.05). Reference comparisons used age/sex-adjusted residuals (Wilcoxon rank-sum, BH-FDR; Table S9), framed as contextual because group is confounded with processing batch. Robustness of the eight long-term EAA findings meeting within-family BH-FDR<0.05 was evaluated by removing each of the 11 participants in turn and recomputing the paired Wilcoxon test and paired-sample Cohen’s d (88 refits total), together with visual inspection of individual TP1 to TP3 trajectories (Table S18; Figure S9).

### Statistics: inferred immune composition

As a global sensitivity test, we evaluated the 11 methylation-inferred cell fractions entered in the primary IAA model (the 12-cell reference with monocytes as the implicit reference). For each cell fraction, the paired TP3-minus-TP1 differences were standardized by their across-participant standard deviation. The global statistic was defined as n multiplied by the sum of the squared mean standardized differences across the 11 fractions. Its exact null distribution was obtained by exhaustively applying all 2^11^=2,048 subject-level sign patterns to the paired 11-fraction difference vectors, preserving the within-participant correlation among fractions. This avoids the covariance-matrix inversion required by Hotelling’s T^2^, which is unstable when the number of pairs and variables are both 11. Cell-specific contributions to T were treated descriptively (Table S19; Figure S10).

### Statistics: EWAS

The software package limma [14] was applied to M-values with within-subject blocking (duplicateCorrelation, block = subject). The acute EWAS included Age, Sex, and control-probe PCs 1-3. For the long-term EWAS, the no-PC model was used as the primary descriptive model because PC1 was aligned with timepoint; a model including PCs 1-3 was reported as a sensitivity analysis. Neither model can uniquely separate technical from biological timepoint variation under complete alignment. A second model added inferred immune-cell proportions. Genomic inflation (lambda) was reported. BACON empirical-null correction [15] was applied because raw BH-FDR is untrustworthy at lambda=1.66 for the long-term contrast (calibration in Figure S3; Table S13). CpGs were reported at suggestive P < 1 x 10^-4^; genome-wide significance at BACON-FDR<0.05 (none survived in either contrast). Over-representation analysis of suggestive CpG lists used clusterProfiler (GO BP/MF/CC, KEGG) [16] and MSigDB Hallmark [17], ranked by nominal p with FDR flagged; direction-split (hyper/hypo) enrichment is in Figure S7. Regions were called with DMRcate (region-level HMFDR; Fisher/Stouffer combined p); DMR-associated genes were themselves submitted to clusterProfiler GO/KEGG over-representation. DMLs, DML pathways, DMRs, and DMR-gene pathways are reported in Tables 5-8 (full lists Tables S5-S8); the combined acute-vs-long-term epigenome-wide figure is Figure 4.

**Fig. 4.**
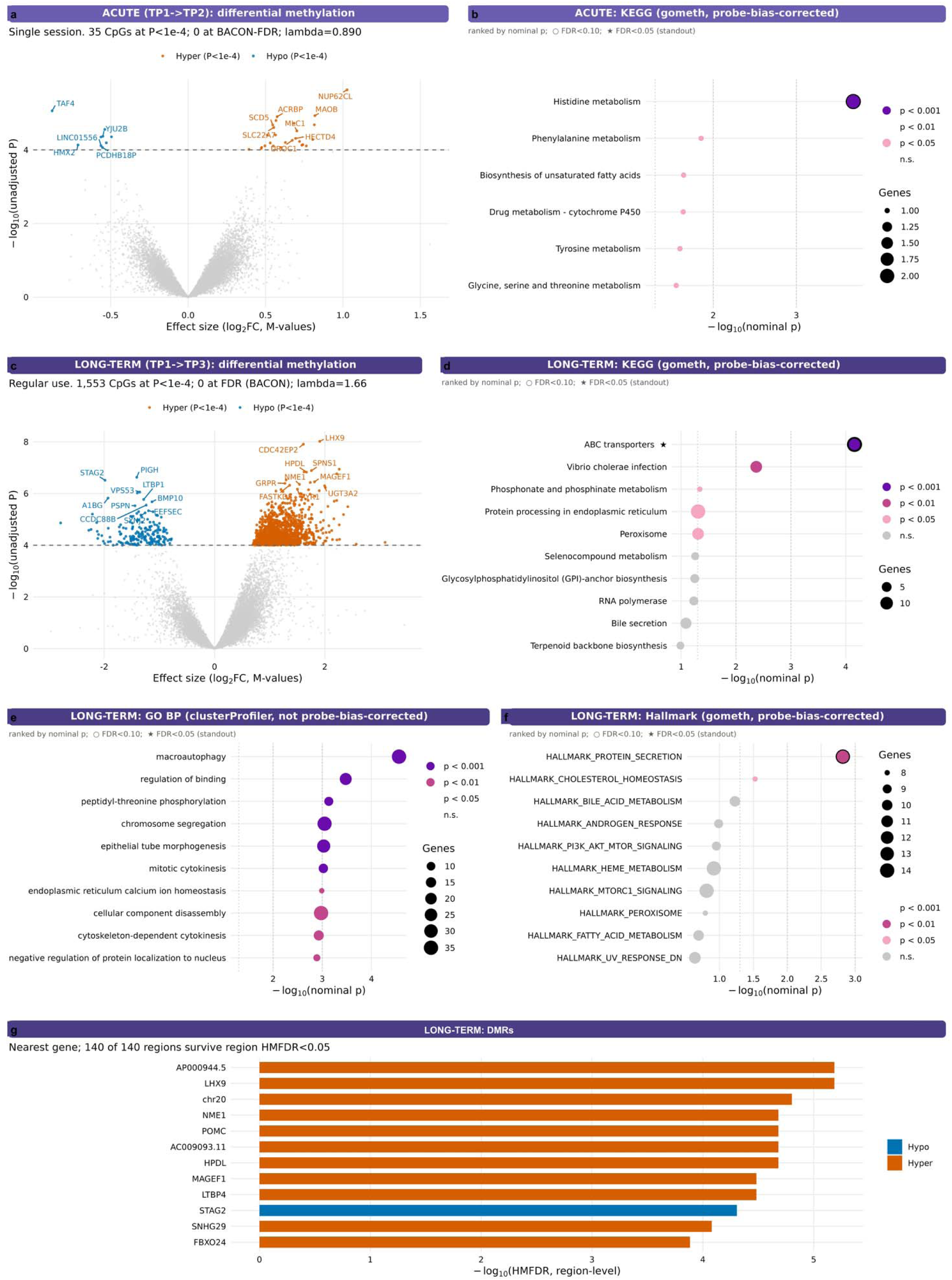
Epigenome-wide analyses of acute and long-term contrasts. Acute TP1–TP2 analyses are shown in row 1: (a) volcano plot from the age-, sex-, and control-probe-PC1–3-adjusted model (35 CpGs at raw P<1×10-4; 126 CpGs at BACON-calibrated p<1×10-4; lambda=0.890; BACON inflation=0.926; no CpG at BACON-FDR<0.05) and (b) probe-bias-corrected gometh KEGG enrichment, for which the leading term was histidine metabolism (p=2.06×10-4, FDR=0.0768). Long-term TP1–TP3 analyses are shown in rows 2–4: (c) volcano plot from the primary no-PC model (1,553 CpGs at raw P<1×10-4; 329 CpGs at BACON-calibrated p<1×10-4; lambda=1.664; BACON inflation=1.072; no CpG at BACON-FDR<0.05); (d) probe-bias-corrected gometh KEGG enrichment, including ABC transporters at FDR=0.0258; (e) secondary clusterProfiler GO Biological Process enrichment; (f) probe-bias-corrected gometh MSigDB Hallmark enrichment; and (g) 12 selected long-term DMRcate regions from the 140 regions meeting HMFDR<0.05. The long-term with-PC sensitivity model identified 360 CpGs at raw P<1×10-4. DMRcate identified 5 acute and 140 long-term regions at HMFDR<0.05; the five acute regions are reported in Table 7 and Supplementary Table S7 but are not displayed in panel g. Source statistics are in Tables 5–7 and Supplementary Tables S5–S7 and S13

**Table 5.**
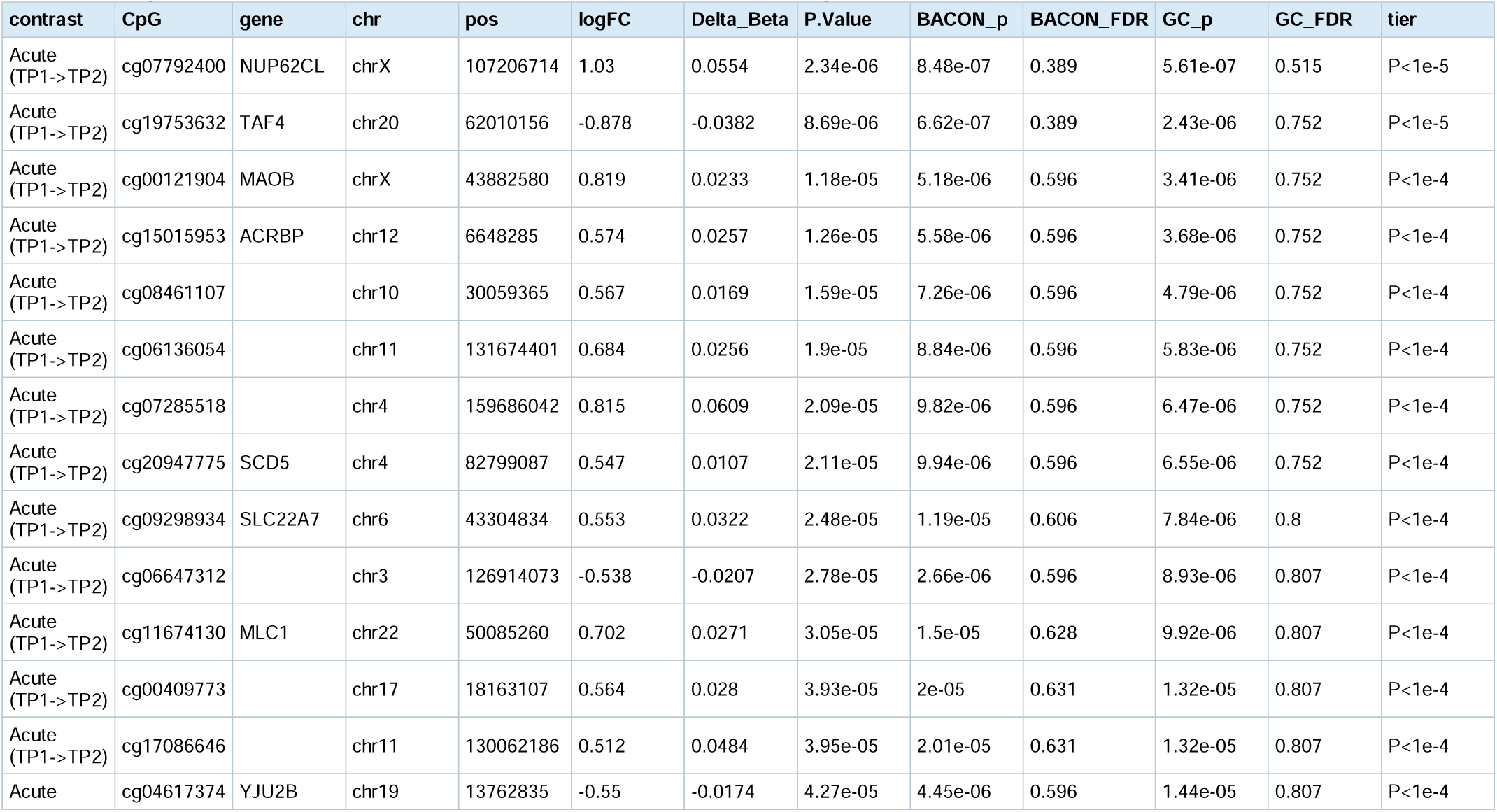
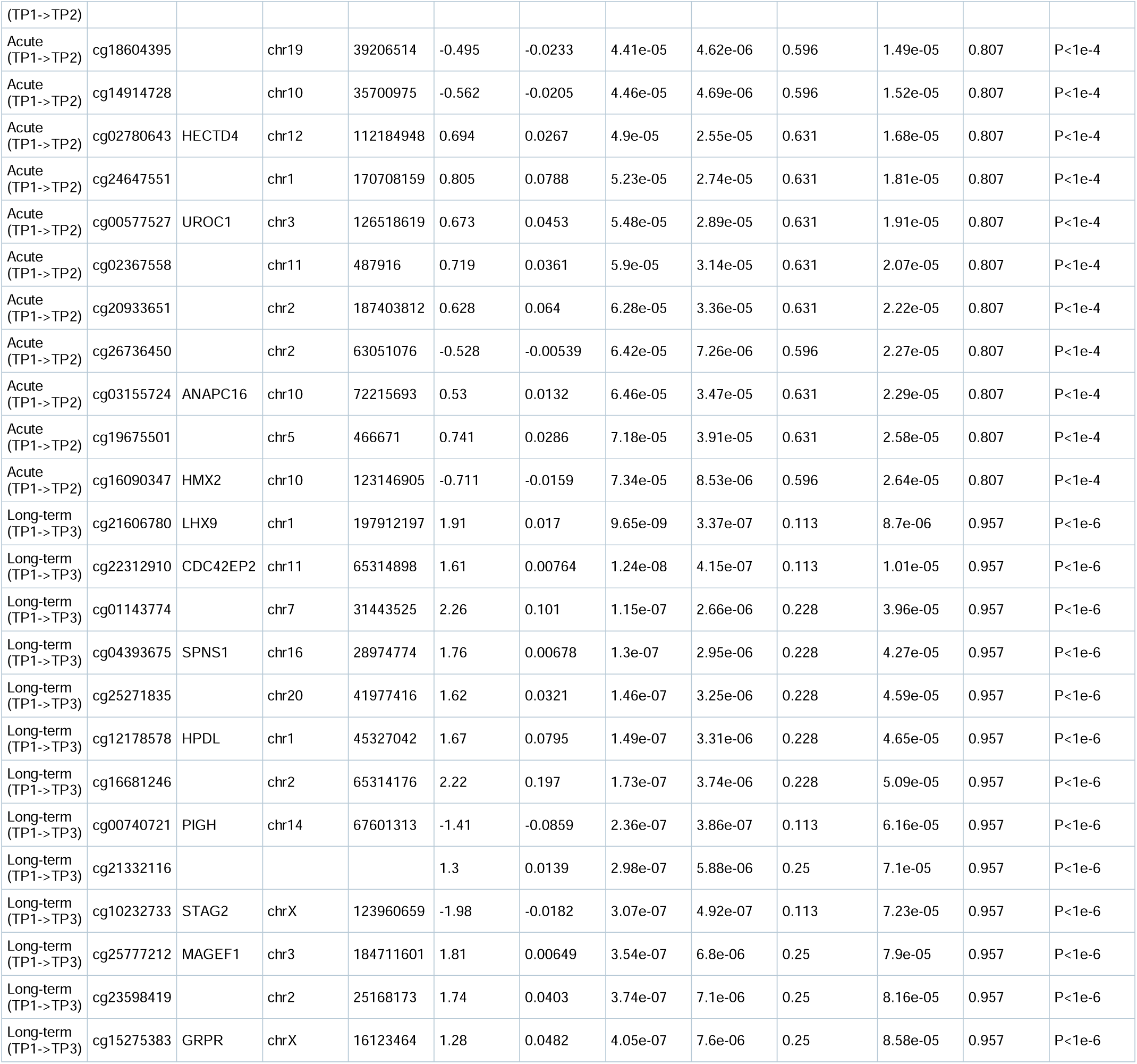
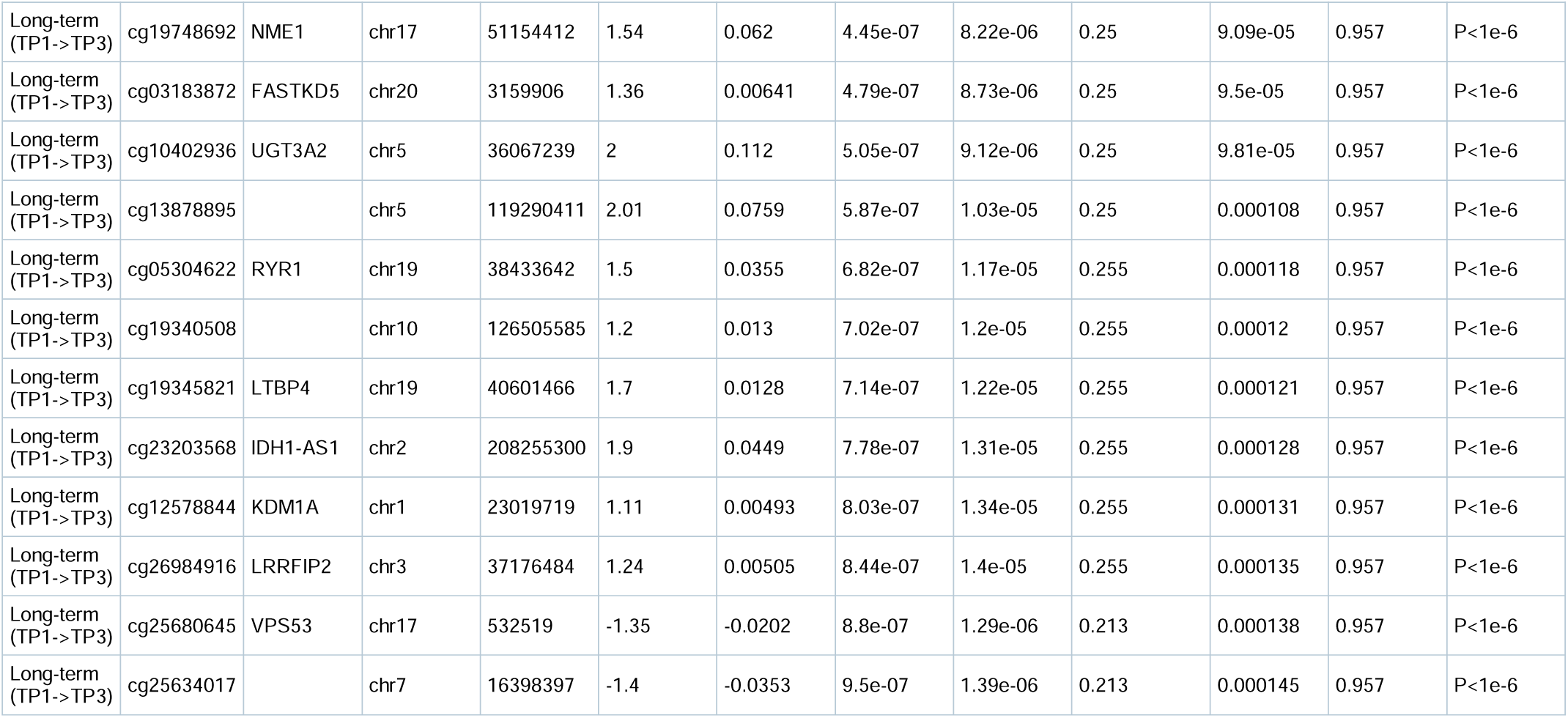
Selected differentially methylated loci. The acute and long-term CpGs with the smallest raw P values among loci meeting raw P<1×10-4 are shown with genomic annotation, log-fold change on M values, mean beta-value difference, raw P, BACON-calibrated p and FDR, genomic-control p and FDR, and reporting tier. The complete raw-P<1×10-4 sets (acute and long-term CpGs) are in Supplementary Table S5; BACON calibration summaries are in Supplementary Table S13.

**Table 6.** Selected pathway annotations of suggestive DMLs. Primary, probe-bias-corrected gometh results are shown: the leading acute KEGG terms, the leading long-term KEGG terms, and the leading long-term MSigDB Hallmark terms. The complete primary gometh and secondary clusterProfiler results are in Supplementary Table S6.

| contrast | collection | method | term | genes | p | FDR | tier |
| --- | --- | --- | --- | --- | --- | --- | --- |
| Long-term (TP1->TP3) | KEGG | gometh (primary, probe-bias-corrected) | ABC transporters | 10 | 6.92e-05 | 0.0258 | FDR<0.05 |
| Long-term (TP1->TP3) | KEGG | gometh (primary, probe-bias-corrected) | Vibrio cholerae infection | 8 | 0.00428 | 0.796 | p<0.01 |
| Long-term (TP1->TP3) | KEGG | gometh (primary, probe-bias-corrected) | Phosphonate and phosphinate metabolism | 2 | 0.0458 | 1 | p<0.05 |
| Long-term (TP1->TP3) | KEGG | gometh (primary, probe-bias-corrected) | Protein processing in endoplasmic reticulum | 14 | 0.0491 | 1 | p<0.05 |
| Long-term (TP1->TP3) | KEGG | gometh (primary, probe-bias-corrected) | Peroxisome | 8 | 0.0492 | 1 | p<0.05 |
| Acute (TP1->TP2) | KEGG | gometh (primary, probe-bias-corrected) | Histidine metabolism | 2 | 0.000206 | 0.0768 | p<0.001 |
| Acute (TP1->TP2) | KEGG | gometh (primary, probe-bias-corrected) | Phenylalanine metabolism | 1 | 0.014 | 1 | p<0.05 |
| Acute (TP1->TP2) | KEGG | gometh (primary, probe-bias-corrected) | Biosynthesis of unsaturated fatty acids | 1 | 0.0227 | 1 | p<0.05 |
| Acute (TP1->TP2) | KEGG | gometh (primary, probe-bias-corrected) | Drug metabolism - cytochrome P450 | 1 | 0.023 | 1 | p<0.05 |
| Acute (TP1->TP2) | KEGG | gometh (primary, probe-bias-corrected) | Tyrosine metabolism | 1 | 0.0252 | 1 | p<0.05 |
| Acute (TP1->TP2) | KEGG | gometh (primary, probe-bias-corrected) | Glycine, serine and threonine metabolism | 1 | 0.0279 | 1 | p<0.05 |
| Acute (TP1->TP2) | KEGG | gometh (primary, probe-bias-corrected) | Tryptophan metabolism | 1 | 0.0319 | 1 | p<0.05 |
| Acute (TP1->TP2) | KEGG | gometh (primary, probe-bias-corrected) | Basal transcription factors | 1 | 0.035 | 1 | p<0.05 |
| Acute (TP1->TP2) | KEGG | gometh (primary, probe-bias-corrected) | Arginine and proline metabolism | 1 | 0.0413 | 1 | p<0.05 |
| Acute (TP1->TP2) | KEGG | gometh (primary, probe-bias-corrected) | Cocaine addiction | 1 | 0.0493 | 1 | p<0.05 |
| Long-term (TP1->TP3) | Hallmark | gometh (primary, probe-bias-corrected) | HALLMARK_PROTEIN_SECRETION | 13 | 0.00152 | 0.076 | p<0.01 |
| Long-term (TP1->TP3) | Hallmark | gometh (primary, probe-bias-corrected) | HALLMARK_CHOLESTEROL_HOMEOSTASIS | 8 | 0.0299 | 0.747 | p<0.05 |

**Table 7.** Selected differentially methylated regions. All acute DMRcate regions and the leading long-term regions meeting HMFDR<0.05 are shown with genomic interval, nearest-gene annotation, CpG count, mean beta-value difference, Fisher combined p value, HMFDR, and reporting tier. The complete set of HMFDR-positive regions (acute and long-term) is in Supplementary Table S7.

| contrast | region | gene | no.cpgs | meandiff | Fisher | HMFDR | tier |
| --- | --- | --- | --- | --- | --- | --- | --- |
| Acute (TP1->TP2) | chr12:6648091-6648285 |  | 3 | 0.0118 | 0.000101 | 8.19e-05 | FDR<0.05 |
| Acute (TP1->TP2) | chrX:107206496-107206955 | DNAAF6 | 11 | 0.0131 | 0.000378 | 8.19e-05 | FDR<0.05 |
| Acute (TP1->TP2) | chr4:159686042-159686180 | LINC02233 | 2 | 0.0722 | 0.000101 | 8.19e-05 | FDR<0.05 |
| Acute (TP1->TP2) | chr20:62010156-62010203 | TAF4 | 2 | -0.0269 | 0.000129 | 8.19e-05 | FDR<0.05 |
| Acute (TP1->TP2) | chr6:28944096-28944389 | LINC01556 | 9 | -0.0205 | 2.55e-05 | 0.00076 | FDR<0.05 |
| Long-term (TP1->TP3) | chr11:65314599-65315321 | AP000944.5 | 6 | 0.00986 | 4.21e-08 | 6.5e-06 | FDR<0.05 |
| Long-term (TP1->TP3) | chr1:197912197-197912371 | LHX9 | 5 | 0.0149 | 1.14e-06 | 6.5e-06 | FDR<0.05 |
| Long-term (TP1->TP3) | chr20:41977416-41977580 |  | 2 | 0.0883 | 2.38e-08 | 1.57e-05 | FDR<0.05 |
| Long-term (TP1->TP3) | chr17:51153868-51154412 | NME1 | 2 | 0.0685 | 1.84e-08 | 2.08e-05 | FDR<0.05 |
| Long-term (TP1->TP3) | chr2:25168173-25168405 | POMC | 2 | 0.0354 | 4.29e-08 | 2.08e-05 | FDR<0.05 |
| Long-term (TP1->TP3) | chr16:28974505-28975032 | AC009093.11 | 8 | 0.00606 | 1.95e-09 | 2.08e-05 | FDR<0.05 |
| Long-term (TP1->TP3) | chr1:45326870-45327360 | HPDL | 7 | 0.0388 | 7.87e-10 | 2.08e-05 | FDR<0.05 |
| Long-term (TP1->TP3) | chr3:184711601-184711995 | MAGEF1 | 6 | 0.00636 | 3.35e-09 | 3.29e-05 | FDR<0.05 |
| Long-term (TP1->TP3) | chr19:40601466-40601667 | LTBP4 | 3 | 0.00719 | 5.49e-05 | 3.29e-05 | FDR<0.05 |
| Long-term (TP1->TP3) | chrX:123960151-123961034 | STAG2 | 12 | -0.0369 | 5.59e-10 | 4.93e-05 | FDR<0.05 |
| Long-term (TP1->TP3) | chr17:16439820-16440085 | SNHG29 | 4 | 0.0422 | 2.07e-08 | 8.32e-05 | FDR<0.05 |
| Long-term (TP1->TP3) | chr7:100585599-100585994 | FBXO24 | 4 | 0.0148 | 5.59e-10 | 0.000131 | FDR<0.05 |
| Long-term (TP1->TP3) | chr5:94618855-94619160 | SLF1 | 3 | 0.0189 | 1.5e-07 | 0.000131 | FDR<0.05 |
| Long-term (TP1->TP3) | chr3:184363019-184363434 | POLR2H | 10 | 0.00769 | 5.11e-08 | 0.000145 | FDR<0.05 |
| Long-term (TP1->TP3) | chr16:69329961-69330117 | COG8 | 2 | 0.0113 | 3.48e-06 | 0.00016 | FDR<0.05 |
| Long-term (TP1->TP3) | chr20:41137145-41137720 | PLCG1 | 9 | 0.00283 | 1.3e-09 | 0.000214 | FDR<0.05 |
| Long-term (TP1->TP3) | chr4:139016269-139016320 | NOCT | 3 | 0.0458 | 2.85e-07 | 0.000214 | FDR<0.05 |
| Long-term (TP1->TP3) | chr14:49598853-49599300 | LRR1 | 4 | 0.0162 | 3.69e-09 | 0.000234 | FDR<0.05 |
| Long-term (TP1->TP3) | chr4:39638421-39639300 | AC108471.2 | 13 | 0.0111 | 1.21e-11 | 0.000296 | FDR<0.05 |
| Long-term (TP1->TP3) | chr1:228082214-228082782 | ARF1 | 13 | 0.00414 | 1.08e-10 | 0.00031 | FDR<0.05 |
| Long-term (TP1->TP3) | chr14:49620693-49620826 | MGAT2 | 4 | 0.00342 | 1.3e-06 | 0.000343 | FDR<0.05 |
| Long-term (TP1->TP3) | chr11:66843121-66843313 | C11orf80 | 7 | 0.00979 | 2.12e-09 | 0.000379 | FDR<0.05 |
| Long-term (TP1->TP3) | chr11:119121326-119121817 | HINFP | 10 | 0.00819 | 1.31e-10 | 0.000499 | FDR<0.05 |
| Long-term (TP1->TP3) | chr22:31945205-31945398 | Z82190.2 | 4 | 0.0128 | 0.000725 | 0.000499 | FDR<0.05 |
| Long-term (TP1->TP3) | chr3:130850190-130850988 | ATP2C1 | 8 | 0.0297 | 1.3e-13 | 0.00055 | FDR<0.05 |

**Table 8.**
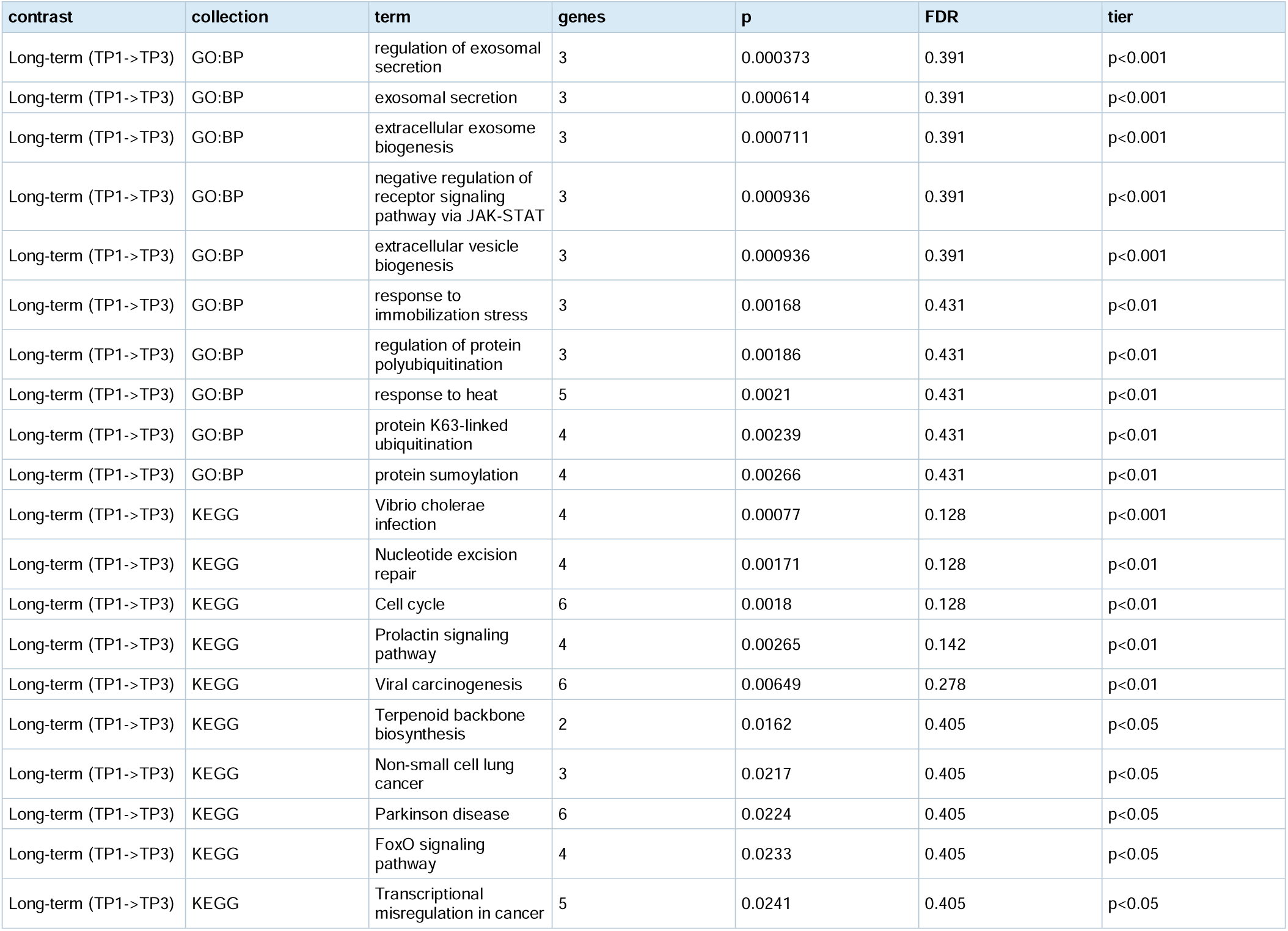
Selected pathway annotations of long-term DMR-associated genes. The leading GO Biological Process terms and KEGG terms derived from genes annotated to the long-term HMFDR-positive regions are shown with gene counts, nominal p values, FDR values, and reporting tiers. Acute DMR-gene enrichment was not performed because only four acute regions had gene annotations. Complete long-term results are in Supplementary Table S8.

## Results

### 2.1 Cohort, design, and tolerability

18 firefighters (15 males and three females) with a mean age of 42.6 years and an age range of 24.9 – 54.8 years provided whole-blood methylation at up to three timepoints (19 enrolled; 18 with methylation microarray data). One participant was excluded from all paired contrasts (non-conforming draw pattern; see Methods), leaving 17 at baseline/time point 1 (TP1), 10 acute/time point 2 (TP2), and 11 long-term/time point three (TP3) (Table 1; Figure 1). Three further subjects’ second draws were not collected during the same session and were excluded from the acute contrast only (see Methods). The resulting paired samples included 10 participants in the acute contrast (all male) and 11 participants in the long-term contrast (10 male and 1 female).

Supporting analyses follow the analytic workflow: quality control and technical calibration (Figures S1-S3), baseline quality of life and adherence (Figures S4-S5), cell-reference and enrichment sensitivity analyses (Figures S6-S7), and technical, subject-level, and global robustness checks (Figures S8-S10). The Supplementary Tables proceed from full predictor panels (Tables S1-S4) through EWAS features and annotations (Tables S5-S8), reference-cohort context (Table S9), inferred-cell analyses (Tables S10-S12), EWAS genomic inflation and BACON empirical null summary (Table S13), quality-of-life and adherence records (Tables S14-S16), and technical and robustness analyses (Tables S17-S19).

Effects were tested as paired within-subject Wilcoxon contrasts for the acute (single session, same-day) and long-term (8-18 weeks of regular use) responses. A total of 188 session-level entries were submitted to the hydration-tracking form. Among the eight long-term participants with individually linkable adherence records, session counts summed to 121 (median 15.5 per participant, range 1-32) at a typical recorded device setting of 6/7 (IQR 5-6). Two mild symptoms were recorded among submitted entries (one headache and one report of dry mouth); incomplete and uneven logging limited inference regarding adherence and tolerability (Figure S5 and Tables S15-S16). A reference cohort of 36 disease-free, age/sex-matched adults (30 males and six females with a mean age of 42.8 years; age-matched, Wilcoxon p=0.847) serves as a contextual comparison (Table 1).

### 2.2 Acute epigenetic response following a single sauna session

A single sauna session produced no epigenetic-age response surviving multiple-testing correction. Across 613 predictors, none survived within-family BH-FDR under either age/sex-adjusted epigenetic age acceleration (EAA) (55/613 at nominal p<0.05) or 12-cell leukocyte-composition-adjusted acceleration (IAA) (13/613 at nominal p<0.05) (Tables 2-4 and Tables S1-S4). The SystemsAge Hormone organ clock and DNAmDHEAS showed nominal decreases after one session: Hormone decreased (d=-1.08, p=0.0098, FDR=0.12) and DNAmDHEAS decreased (d=-0.78, p=0.020, FDR=0.20); neither survives correction across the full predictor family. DNAmDHEAS moved in the opposite direction long term, whereas the long-term Systems Age Hormone score did not reach nominal significance (d=+0.60, p=0.083). DunedinPACE also decreased at the nominal level on EAA (d=-0.77, p=0.027, FDR=0.554), though not on IAA (d=-0.67, p=0.065, FDR=0.831, not nominally significant). This was the same direction as, not opposite to, the long-term trend (Section 2.3); 20 of the 34 clocks in this family move numerically in the decreasing direction after a single session, and of these, DunedinPACE and HbA1c-MRS individually reach nominal significance. The single strongest nominal hit in the whole epigenetic clocks family, however, moves the opposite way: PCPAI1 increased (d=+1.14, p=0.0039, FDR=0.13), a GrimAge-component score, though like the others it does not survive within-family correction. These nominal acute findings identify candidate transient responses but do not establish concrete benefit, adaptation, or hormesis. Monocyte proportion showed a similar nominal decrease (d=-0.83, p=0.020, FDR=0.21); no other 12-cell immune proportion reached nominal significance (Table S10).

At the epigenome-wide level, the acute contrast yielded 35 CpGs at P < 1 x 10^-4^ (model includes control-probe PC1-3; lambda=0.890, BACON inflation=0.926), below the approximately 92 expected by chance across 918,757 tested sites, with 0 CpGs at BACON-FDR<0.05. BACON calibration nevertheless identified 126 CpGs at p_bacon<0.0001, and DMRcate returned 5 acute regions at HMFDR<0.05, providing focused candidates for investigation of the acute response (Section 2.10). No CpG reached the corrected threshold (Table 5 and Table S5). No pathway survived probe-bias-corrected enrichment: under missMethyl *gometh*, the strongest KEGG term (histidine metabolism) reached only FDR=0.077 (nominal P=2.1 x 10^-4) and MSigDB Hallmark showed no signal at all (FDR=1 throughout). A clusterProfiler over-representation test flagged inner-ear morphogenesis and related Gene Ontology (GO) terms at nominal FDR, but as with the long-term analysis (see below starting at Section 2.8), over-representation testing does not correct for the number of CpGs per gene and is not supported by the probe-bias-corrected method; it is not reported as a finding (Table 6 and Table S6). The acute single-session analysis therefore identified a focused set of candidate CpGs and regions without within-family-corrected predictor or FDR-significant pathway changes, with nominal observations reported explicitly as nominal and hypothesis-generating. Per protocol, the acute (TP2) draw was collected within about an hour of the first sauna session, a gap consistent with published acute sauna leukocyte kinetics in which even the slowest-clearing leukocyte fraction returns to near baseline within 30 minutes of a single Finnish sauna session [18], and with sauna’s immunomodulatory literature more broadly, in which a measurable immune-parameter change requires a series of sessions rather than one exposure [19].

### 2.3 Long-term follow-up shows a mixed timepoint-associated predictor pattern (TP1 to TP3)

On age/sex-adjusted EAA, the long-term contrast (n=11) produced within-family BH-FDR-significant change in eight predictors spanning four classes (Table 2 and Table S1; Figure 2a–b): decreased PCDNAmTL (d=-1.92, FDR=0.040); an increased depression-risk methylation score, DepressionBarbu (d=+1.75, FDR=0.040); increased SystemsAge Blood age (d=+1.20, FDR=0.029) and SystemsAge Lung age (d=+1.02, FDR=0.029); and increased DNAm PhysAge subcomponents (Table 3 and Table S2): DNAmPulsePressure (d=+1.35, FDR=0.023), DNAmCystatinC (d=+1.05, FDR=0.023), and DNAmDHEAS increased (d=+0.92, FDR=0.034). However, DNAmHDL decreased (d=-1.08, FDR=0.023).

All eight findings were robust to leave-one-participant-out resampling: across 88 refits involving 11 participants and eight predictors, no participant’s removal reversed an effect direction or raised the nominal Wilcoxon p value to >=0.05 (largest leave-one-out p=0.0273; Table S18). Individual trajectories likewise show that the paired changes were not attributable to any single extreme participant (Figure S9). This analysis addresses single-participant influence but does not resolve the timepoint/chip confounding described below.

A substantially larger cluster reached nominal significance (p=0.005-0.010) but landed just above the within-family threshold once the complete clocks family was tested (83 candidate predictors, of which 81 yielded analyzable paired tests and entered the FDR correction; FDR=0.057 for all of them, versus FDR<0.05 under the previous, incomplete 35-member family): the aging-rate/mitotic group pcgtAge (d=+1.17), irS and irS2 (d=+1.01), and tnsc2/tnsc (mitotic-rate variants, d=+1.01), together with AdaptAge (d=+1.00), ENCen100 (d=+1.02), Retroclockv2 (d=-1.03), RepliTaliNorm (d=-0.98), LeeRefinedRobust (d=-0.98), XRa (d=-0.81), and PCLeptin (d=-1.09). Liver (d=+0.78, p=0.024, FDR=0.098), PCHannum (d=+0.82, p=0.024, FDR=0.11), Retroclock (d=-0.91, p=0.014, FDR=0.069), and DNAmClockCortical (d=+0.99, p=0.014, FDR=0.069) reached nominal but weaker significance. The SystemsAge Hormone age (d=+0.60, p=0.083), Brain age (d=+0.75, p=0.067), and Metabolic age (d=+0.77, p=0.054) clocks did not reach even nominal significance. HbA1c MRS and the Marioni type-2-diabetes EpiScore remained non-significant under both EAA and IAA.

Critically, several next-generation epigenetic aging clocks [20] did not change. PCGrimAge (d=-0.23, p=0.52), PhenoAge (d=+0.40, p=0.32), and OMICmAge (d=-0.11, p=0.70) were all flat. PCPhenoAge (a principal component-clock variant of PhenoAge) showed a nominal, not FDR-significant, increase on EAA (d=+0.71, p=0.042, FDR=0.16) that was attenuated after leukocyte-composition adjustment (IAA d=-0.30, p=0.70). DunedinPACE (pace-of-aging) trended favorably (decreasing) but did not reach nominal significance in this contrast (EAA d=-0.67, p=0.102; IAA d=-0.54, p=0.102), unlike the acute contrast (Section 2.2), where the same predictor did reach nominal significance. Thus the FDR-significant long-term signal was confined to telomere, behavioral/psychiatric, organ-system, and metabolic-proxy predictors, with the aging-rate/mitotic cluster reaching only nominal significance and no within-family FDR-confirmed change in the mortality/pace-predictive clocks. These heterogeneous methylation-score changes should not be interpreted as direct changes in organ age or physiology.

### 2.4 Long-term predictor associations are sensitive to leukocyte-composition adjustment

Adjusting for 11 immune-cell deconvolution reference (IAA; Age+Sex+cells) attenuated every one of the eight EAA-FDR-significant findings mentioned above to non-significance: PCDNAmTL (EAA FDR=0.040; IAA d=-0.03, p=0.70, FDR=0.96), DepressionBarbu (EAA FDR=0.040; IAA d=+0.96, p=0.024, FDR=0.33; nominal only), SystemsAge Blood age (EAA FDR=0.029; IAA d=+0.25, p=0.577, FDR=0.77), SystemsAge Lung age (EAA FDR=0.029; IAA d=+0.08, p=0.577, FDR=0.77), DNAmPulsePressure (EAA FDR=0.023; IAA d=+0.03, p=0.765, FDR=1.0), DNAmHDL (EAA FDR=0.023; IAA d=-0.55, p=0.102, FDR=0.52), DNAmCystatinC (EAA FDR=0.023; IAA d=+0.35, p=0.206, FDR=0.52), and DNAmDHEAS (EAA FDR=0.034; IAA d=+0.41, p=0.206, FDR=0.52). Zero predictors in the confirmatory Clocks/SystemsAge/PhysAge panel reach IAA-FDR<0.05 (Figure 2c; Tables 2-3). Unlike the previous, incomplete predictor panel, there is no confirmed exception in the corrected accounting. The one immune-adjacent SystemsAge organ clock, Inflammation, was flat under both EAA (p=0.898) and IAA (p=0.278). These results show that inferential status is sensitive to adjustment for methylation-inferred leukocyte composition.

PCLeptin, the candidate least attenuated by this cell-composition adjustment in the previous predictor panel, no longer reaches EAA-FDR<0.05 once the panel was completed (EAA d=-1.09, p=0.00977, FDR=0.057; nominal, not within-family significant); its IAA effect remains of comparable magnitude (d=-1.12, p=0.00977, FDR=0.329), so the qualitative pattern of a signal retained after this adjustment persists, but only as a nominal, model-dependent observation. Four predictors show a pattern compatible with statistical suppression, with null or near-null results on EAA and nominal significance only after adjustment. None survived within-family correction: PCADM (a GrimAge mortality-component score; EAA d=-0.67, p=0.054; IAA d=-0.91, p=0.019, FDR=0.33), PCCystatinC (EAA d=-0.44, p=0.465, null; IAA d=-0.75, p=0.024, FDR=0.33), and, newly visible once the panel was completed, DamAge (EAA d=-0.62, p=0.102, null; IAA d=-0.90, p=0.024, FDR=0.33) and Stochastic.PhenoAge (EAA d=-0.43, p=0.24, null; IAA d=-0.76, p=0.032, FDR=0.37). We flag all four as hypothesis-generating, not confirmed. A claim that adjustment specifically unmasks a favorable biological signal is not supported because none reaches within-family significance. A 19-cell CAB-refined deconvolution sensitivity analysis shows a broadly consistent picture, with one notable divergence: DNAmCystatinC reaches IAA-FDR<0.05 under the 19-cell reference (d=+0.82, p=0.000977, FDR=0.0098) but not under the primary 12-cell reference (FDR=0.52). All 11 subjects shifted in the same direction under the 19-cell model; however, because statistical significance depended on the deconvolution reference, the result is reported as a sensitivity-dependent observation and is not adopted as a primary finding, per this study’s established convention that the 19-cell reference is a secondary consistency check (Figure S6; Table S12). pcgtAge, irS, irS2, tnsc2/tnsc, and AdaptAge were nominally significant under both references but were not FDR-significant under either. These results were directionally consistent with, but not identical to, the 12-cell result.

### 2.5 Separately processed reference-cohort comparison

Compared with age/sex-matched disease-free controls (age/sex-adjusted EAA, 103-predictor comparison panel, which was expanded from 32 previously tested predictors once this comparison’s predictor list was cross-checked against the corrected Clocks family), firefighters at baseline showed ten predictors at FDR<0.05 (Table S9; Figure 3a), a mixed picture rather than a uniform direction: lower values for pcgtAge (d=-0.85, FDR=0.041), irS (d=-0.82, FDR=0.043), irS2 (d=-0.84, FDR=0.043), tnsc2/tnsc (d=-0.81, FDR=0.043), and AdaptAge (d=-1.20, FDR=0.024); higher predicted cardiorespiratory fitness on DNAmVO_2_max (d=+1.40, FDR=0.0093); lower ENCen100 (d=-0.94, FDR=0.023); and, in the opposite direction, higher predicted waist-to-hip ratio on DNAmWHR (d=+0.93, FDR=0.024) and higher DamAge (d=+0.97, FDR=0.041). DunedinPACE showed a nominal, not FDR-significant, difference in the higher direction (d=+0.81, p=0.015, FDR=0.078; unchanged from the previous panel, since DunedinPACE was already tested). At post-program follow-up, three predictors reached FDR<0.05 (Table S9; Figure 3a): higher predicted cardiorespiratory fitness on DNAmVO_2_max (d=+2.32, FDR=0.00067, strengthened relative to baseline), higher DepressionBarbu (d=+1.14, FDR=0.039), and lower hypoSC (d=-0.95, FDR=0.039). Because controls were processed in a separate run (group perfectly confounded with processing batch; Figure S2), this comparison is descriptive and non-confirmatory. It is not used to infer an intervention effect.

### 2.6 Inferred leukocyte composition

Individual immune proportions showed only a nominal Treg increase (d=+0.72, p=0.032, FDR=0.386), with nothing surviving FDR (Table S11; Figure 3b). No specific leukocyte population was confirmed, and the analysis does not estimate a formal mediated or indirect effect. A global exact sign-flip test of the 11 cell fractions used in the IAA model was directionally suggestive but not significant (T=18.788, exact p=0.0967; 2,048 sign patterns; Table S19; Figure S10). Treg and Bmem contributed 30.5% and 21.0% of the statistic, respectively.

### 2.7 Epigenetic Biomarker Proxy (EBP) metabolite proxies and Marioni protein EpiScores

Recomputed on clean age/sex EAA, the EBP v2 metabolite-proxy panel (1,378 scores tested in the long-term contrast) showed 99 at within-panel FDR<0.05 on EAA; none survived 12-cell IAA (0/1,378 IAA-FDR<0.05) (Table 4; full Table S3). This was a large age/sex-adjusted signal whose inferential status is highly sensitive to leukocyte-composition adjustment. Among the 113 Marioni protein EpiScores tested in the long-term contrast (Table 4; full Table S4), 17 reached even nominal significance on EAA or IAA, and none reached within-family BH-FDR<0.05 under either adjustment. The panel was expanded from the 69 previously tested scores after its dedicated pipeline output was cross-checked for completeness. This panel is accordingly reported as exploratory throughout, with no confirmed individual protein-level finding. The strongest nominal leads (uncorrected, hypothesis-generating only) were BMP-1 (EAA d=+1.05, p=0.00098; IAA p=0.28), MIA (EAA d=+1.31, p=0.0020; IAA d=+0.74, p=0.032), Testican-2 (EAA d=-1.30, p=0.0020; IAA p=0.12), Osteomodulin (EAA d=+1.05, p=0.0098; IAA d=+1.06, p=0.0098), and GHR (EAA d=+1.09, p=0.0068; IAA d=+0.60, p=0.083). The smaller EBP v1 panel (397 scores tested in the long-term contrast, used to decompose OMICmAge, which was itself flat) showed the same overall pattern at a larger scale: 79 scores reached within-panel FDR<0.05 on EAA, whereas none survived 12-cell IAA; individual EBP v1 scores are not separately tabulated.

### 2.8 Acute analysis identifies a BACON-calibrated candidate CpG set (TP1 to TP2)

The acute contrast yielded 35 CpGs at the suggestive P less than 0.0001 tier (model includes control-probe PC1 through PC3; lambda = 0.890), below the approximately 92 expected by chance across 918,757 tested sites. BACON empirical-null correction (bacon inflation = 0.926, bias = 0.159) identified 126 CpGs at p_bacon<0.0001, including all 35 raw-P<0.0001 CpGs (Table S13); thus, the BACON-defined suggestive set was larger and was not identical to the raw-P tier. However, 0 CpGs reach BACON-FDR below 0.05 once corrected for the approximately 918,757 tests performed. The BACON inflation estimate of 0.926 provides no evidence of test-statistic inflation in this contrast. The 126 CpGs meeting p_bacon<0.0001 therefore provide a calibrated candidate set for studying acute methylation responses, even though the smaller raw-P tier fell below chance expectation at this sample size.

Under gometh, the primary, probe-bias-corrected method, run for KEGG and Hallmark only, the strongest KEGG term was histidine metabolism (nominal P = 0.00021, FDR = 0.077, not significant); MSigDB Hallmark showed no signal at all, with FDR equal to 1 throughout (Table 6; full results in Table S6). A clusterProfiler GO over-representation test, the secondary, not probe-bias-corrected method, flagged inner-ear morphogenesis and related terms at nominal FDR, but this method does not correct for the number of CpGs per gene and is not reported as a finding. Taken together, the acute DML analysis identified a BACON-calibrated candidate set for follow-up, although it did not yield an FDR-significant gometh pathway.

### 2.9 Long-term analysis identifies BACON-calibrated CpGs and ABC transporter enrichment (TP1 to TP3)

The long-term contrast yielded 1,553 CpGs at the suggestive P less than 0.0001 tier under the primary, no-PC model (a with-PC sensitivity model gave 360), against a genomic-inflation caveat (lambda = 1.664; BACON’s own empirical-null inflation estimate = 1.072, bias = 0.270). After BACON calibration, 329 of the 1,553 raw-P<0.0001 CpGs also met p_bacon<0.0001 (21%). The reduced overlap shows that membership in the suggestive tier is sensitive to empirical-null calibration; it does not by itself classify the remaining loci as false positives. After Benjamini-Hochberg FDR is applied on top of p_bacon across all 918,757 tested sites, 0 CpGs reach BACON-FDR below 0.05 (Table S13), including the 329 that survived the inflation-correction step alone. The strongest loci by raw P-value were *LHX9*, *CDC42EP2*, *SPNS1*, *HPDL*, *PIGH*, and *STAG2* (Table 5; full results in Table S5).

Under gometh, run for KEGG and Hallmark only, the KEGG ABC transporters term reached FDR significance: 10 of 45 pathway genes were represented, with nominal P=0.000069 and FDR=0.026. This probe-bias-corrected result identifies membrane transport as a prominent biological theme among the long-term methylation candidates. Hallmark reached no term below FDR 0.05 (the best was PROTEIN_SECRETION, nominal P = 0.0015, FDR = 0.076) (Table 6; full results in Table S6). A clusterProfiler GO Molecular Function over-representation test, the secondary method, showed a concordant transporter-related theme: ABC-type transporter activity, ATPase-coupled transmembrane transporter activity, and primary active transmembrane transporter activity are among nine GO Molecular Function terms reaching FDR below 0.05 under that method (full list in Table S6). Although clusterProfiler does not correct for unequal CpG coverage across genes, its convergence with the probe-bias-corrected gometh result strengthens the prioritization of membrane-transport biology for follow-up. The long-term DML analysis therefore identified one FDR-significant, probe-bias-corrected KEGG enrichment (ABC transporters), echoed by concordant GO terms from a second method. Together, these results provide a coherent membrane-transport theme for mechanistic follow-up, while the remaining pathway signals are retained as exploratory candidates.

### 2.10 Acute analysis identifies five HMFDR-significant regions (TP1 to TP2)

Region-level analysis using DMRcate returned 5 regions at HMFDR below 0.05 (Table 7; full results in Table S7). The CpG seed set was focusd – 32 of 892,319 probes at raw P<0.0001 – but spatial aggregation identified statistically supported regional signals despite the modest acute sample. These five regions therefore provide specific candidates for investigating early methylation responses to a single portable-sauna session.

Pathway enrichment was not performed: four regions were annotated to *DNAAF6*, *LINC02233*, *TAF4*, and *LINC01556*, and one was intergenic. Four genes are too few for a meaningful enrichment test; accordingly, Table 8 and Table S8 contain no acute rows.

### 2.11 Long-term analysis identifies a pronounced HMFDR-significant regional signal (TP1 to TP3)

Region-level analysis using DMRcate found 140 regions at HMFDR below 0.05, overwhelmingly hypermethylated (134 hyper, 6 hypo), directionally concordant with the CpG-level result above. The strongest regions annotated to *AP000944.5*, *LHX9*, *NME1*, *POMC*, *AC009093.11*, *HPDL*, *MAGEF1*, *LTBP4*, and *STAG2* (Table 7; full results in Table S7). Two of these genes, *LHX9* and *HPDL*, are also among the top CpG-level long-term DML hits described above. The gene *HPDL* is interesting given that reported to guard against oxidative stress and regulate mitochondrial bioenergetics [21].

Analysis of the DMR-associated genes (Table 8; full results in Table S8) did not converge on a coherent biological theme. The leading nominal terms spanned exosomal secretion, JAK-STAT signaling regulation, and protein ubiquitination under GO Biological Process, alongside several unrelated KEGG pathways including Vibrio cholerae infection, nucleotide excision repair, and cell cycle. No GO or KEGG term survived enrichment FDR below 0.05 (the best GO Biological Process FDR was 0.39; the best KEGG FDR was 0.13).

The 140 regions meeting HMFDR<0.05 represent a substantial long-term regional methylation signal and provide a prioritized set of candidate loci for replication and functional study. Although DMR-associated-gene enrichment did not yield an FDR-significant GO or KEGG term, DML-based gometh identified FDR-significant enrichment of KEGG ABC transporters (FDR=0.026), providing an interpretable biological theme for further investigation.

## Discussion

This novel work represents the first investigation of sauna bathing on both clock and proxy epigenetic biomarkers. While exploratory in nature, many of the findings generated here are quite interesting. At baseline, for instance, firefighters were predicted (based on DNA methylation) to have a higher VO_2_ max than the compared contextual cohort. This finding makes intuitive sense as firefighting requires robust physical fitness and the International Association of Firefighters has recommended that firefighters meet a specified VO_2_ max threshold [22]. Firefighters also displayed a higher DamAge and a lower AdaptAge. These two clocks are spinouts from the next-generation epigenetic aging clock CausAge [23] and, as their names imply, AdaptAge is thought to reflect beneficial adaptations while DamAge was trained to quantify accumulated epigenetic damage. Thus, the theoretical ideal is for AdaptAge to be higher and DamAge to be lower. It is intriguing that the opposite pattern is observed here and this could be reflective of the high amounts of physiological stress imposed by this hazardous profession. Indeed, sudden cardiac death has been documented as a major cause of death during the line of duty in United States firefighters [24].

Different epigenetic aging biomarkers provide differential insights and, as such, it is typical to see directionally mixed results [25]. Here, we see that the clock and proxy findings were directionally varied, but informative: eight predictors met within-family BH-FDR<0.05 under age/sex-adjusted EAA after 8-18 weeks of portable-sauna use. DNAmDHEAS increased and PCDNAmTL, a methylation-based telomere-length estimator rather than a direct telomere measurement, decreased under age/sex adjustment but not under the primary 12-cell IAA model. DNAmHDL also decreased, while methylation-derived estimates of depression risk, blood organ age, lung organ age, pulse pressure, and Cystatin-C increased. The broad attenuation of predictor findings under the primary 12-cell adjustment supports immune composition as a contributor and motivates direct immune phenotyping in future studies, even though the corrected cell-specific tests were not conclusive.

Further research is warranted to see if, after a longer time course, these changes resolve or even directionally reverse. These results are reminiscent of what has been previously published for high-intensity interval training, another form of hormetic stress. Prior research looking at vastus lateralis muscle samples has shown that the epigenetic clocks CheekAge, Horvath 2013, Hannum 2013, and Horvath 2018 are all elevated after four weeks in healthy men that underwent high-intensity interval training [26]. Since exercise induces positive inflammatory changes [27] and aging clocks are sensitive to both positive and negative transient stressors [28], these results were interpreted as being reflective of immunological remodeling [26]. It is also germane to mention that other therapeutic interventions, including decitabine, plasmapheresis, and FTC-tenofovir-disoproxil fumarate have been reported to accelerate at least one next-generation epigenetic aging clock [29]. Work in mice has also shown that intermittent hypoxia induces epigenetic age acceleration that reverses after transitioning back to a normoxic environment [30]. Thus, one plausible explanation for the work presented here is that repeated portable-sauna use elicited epigenetic remodeling that includes an immune-composition component.

The clock and methylation-inferred immune analyses together nominate immune composition as a testable component of the long-term response. The inferred-cell changes showed a coordinated pattern, with Treg providing the strongest individual signal (p=0.032, FDR=0.386) and the global sign-flip test approaching significance (p=0.0967). Together with the broad attenuation of EAA findings after cell adjustment, this convergence supports immune composition as a priority for direct phenotyping in subsequent studies. Higher-resolution or sorted-cell methylation could identify the cell populations contributing to this response. Notably, the SystemsAge Inflammation organ clock remained stable under both EAA and IAA, suggesting that the observed pattern may reflect compositional remodeling rather than a generalized inflammatory-aging response.

At the regional level, the long-term contrast produced a substantially broader signal than the acute contrast: 140 versus 5 DMRcate regions met HMFDR<0.05, and 134 of the 140 long-term regions were hypermethylated. At the individual-CpG level, 329 long-term CpGs met the BACON-calibrated, unadjusted threshold p_BACON<1×10^-4^. Together, these findings support a broader epigenetic response after 8–18 weeks of repeated portable-sauna use than after a single session and provide a focused set of high-priority loci for further investigation. Heat-associated molecular remodeling is biologically plausible and supported by prior human studies describing acute blood-mononuclear-cell transcriptional responses to sauna [31], ambient-temperature-associated blood DMRs across exposure windows [32], and time-dependent blood methylation programs after exertional heat illness [33]. Among the long-term DMR annotations identified here, *FOXO3* (7 CpGs; mean Δβ=+0.00885; HMFDR=0.00152) is particularly relevant because HSF1 activates FOXO3 after heat shock in human cells [34]. *HPDL* (7 CpGs; mean Δβ=+0.0388; HMFDR=2.08×10^-5^) is also notable for reported roles in mitochondrial bioenergetics and oxidative-stress protection [21], although a direct role in the heat response has not been demonstrated. These regions therefore represent prioritized candidates for replication, expression studies, and functional assessment of the human response to repeated heat exposure.

This exploratory pilot has several limitations. The acute and long-term contrasts included 10 and 11 paired participants, respectively, there was no thermoneutral control arm, and long-term timepoint was completely aligned with BeadChip, limiting separation of biological and technical timepoint effects. Given the sample size and approximately 919,000 CpGs tested, no individual CpG survived genome-wide FDR correction, although BACON calibration retained 126 acute and 329 long-term CpGs at p_BACON<1×10^-4^ and DMRcate identified five acute and 140 long-term regions at HMFDR<0.05. These constraints limit causal inference, but the internally consistent regional, CpG-level, and pathway signals provide a strong candidate map for replication in larger, controlled studies. Session logs provide limited implementation context for the intervention. Among the eight of 11 long-term participants with individually linkable adherence records, 121 sessions were documented (median 15.5 sessions per participant, range 1–32), while only two mild symptom reports were recorded across 188 submitted entries. These observations are consistent with repeated portable-sauna use and few reported symptoms among the submitted logs. However, incomplete and uneven logging precludes estimation of adherence or tolerability across the full cohort and limits exposure-response analyses. Future studies should use standardized prospective session capture, systematic adverse-event monitoring, and calibrated measures of heat exposure, including chamber temperature and physiological temperature responses.

Based on our findings, more substantial follow-up is more than warranted. A recommended subsequent study would involve a larger number of individuals and a paired cohort that engages in a control activity (such as sitting in thermoneutral environment) at the same time and under the same frequency. Extending the duration of the study and incorporating additional timepoints would also be quite valuable. Additional research looking at other forms of hormetic stress, such as cold exposure, is also justified.

## Abbreviations

BH: Benjamini-Hochberg;
DNAm: DNA methylation;
DMR: differentially methylated region;
EAA: epigenetic age acceleration;
EWAS: epigenome-wide association study;
FDR: false discovery rate; GO, Gene Ontology;
KEGG: Kyoto Encyclopedia of Genes and Genomes;
IAA: leukocyte-composition-adjusted acceleration;
PC: principal component;
TP: timepoint.

## Author contributions

V.B.D.: Conceptualization, methodology, formal analysis, and writing (original draft). A.A.J.: Review, writing, and editing. K.S.: Formal analysis, code review, writing and review. N.K: Patient recruitment, intervention design, study concept. D.S.: TruDiagnostic data organization. S.M.: Study concept and provision of the portable-sauna devices and program logistics. R.S.: Supervision, methodology, and writing (review and editing).

## Acknowledgments

The authors thank the participating firefighters.

## Statements and Declarations

### Ethical Statement

The retrospective secondary analysis of previously collected samples and data reported here was reviewed and approved by the Institute for Regenerative and Cellular Medicine IRB under “Evaluation of TruDiagnostic Internal Dataset” (protocol TRU-012-2022; IRB approval IRCM-2023-269). This retrospective approval did not constitute prospective registration of the intervention. The research was conducted in accordance with the Declaration of Helsinki.

### Consent

Written informed consent was obtained from all participants before participation. The consent described the sauna procedures, three blood collections for epigenetic age and biomarker analysis, research use of de-identified data, and publication of results with participant identities kept confidential.

### Trial registration

Participants were prospectively enrolled in a multi-week portable-sauna intervention. The study was not prospectively registered in a public clinical trial registry. The DNA-methylation analyses reported here were retrospective and exploratory.

## Data availability

Result tables are provided in the Supplementary files. DNA methylation data are available from the corresponding authors (V. Dwaraka and R. Smith) upon request, contingent on the signing of a Data Use Agreement. Interested parties should contact the corresponding authors directly by email to initiate this process. The Data Use Agreement specifies the permitted scope, purpose, and manner of data use, and is required to protect the privacy of study participants. Analysis code will be deposited on Zenodo and made publicly available upon publication of this study.

## AI-tool disclosure

Large-language-model tools assisted with manuscript editing and pre-submission review. Statistical results were generated independently by the study authors and verified against source tables by the authors, who take full responsibility for the accuracy, originality, drafting, and interpretation of the manuscript. Initial draft was written by the authors as well.

## Online Resource 1: Supplementary Figure Legends

**Fig. S1** Post-normalization DNA-methylation quality control. (a) Per-sample ssNoob-normalized beta-value density distributions, colored by timepoint where available. (b) Per-sample medians and interquartile ranges, ordered by timepoint and median. The expected bimodal methylation distributions and similar central tendencies were retained across samples

**Fig. S2** Alignment of control-probe PC1 with long-term timepoint. (a) Paired control-probe PC1 values for the 11 TP1–TP3 pairs; every participant shifted in the positive direction (mean TP3-minus-TP1 change, +0.28). (b) Paired changes for control-probe PCs 1–3. PC1 was uniformly positive, whereas PC2 and PC3 included changes on both sides of zero. This alignment motivated use of the no-PC long-term EWAS as the primary descriptive model and the with-PC model as a sensitivity analysis

**Fig. S3** EWAS calibration and genome-wide view. Quantile–quantile plots show raw, genomic-control-corrected, and BACON-calibrated p values for (a) the acute TP1–TP2 model (raw lambda=0.890; BACON inflation=0.926) and (b) the primary long-term TP1–TP3 model (raw lambda=1.664; BACON inflation=1.072). (c) Manhattan plot of raw long-term p values across the autosomes and sex chromosomes; the dashed line marks raw P=1×10-4. BACON calibration retained 126 acute and 329 long-term CpGs at p<1×10-4; no CpG met BACON-FDR<0.05. Calibration counts are in Supplementary Table S13

**Fig. S4** Baseline WHO quality-of-life item ratings. Means and standard deviations are shown for WHO-QoL items scored on a 1–5 Likert scale; the dashed vertical line marks the midpoint of 3. Item-specific sample sizes were 15 or 16 respondents. These cross-sectional questionnaire data were not linked to the methylation analyses. Source values are in Supplementary Table S14

**Fig. S5** Adherence and tolerability from submitted session logs. (a) Participant-linked session counts for 8 of the 11 long-term participants (121 linked sessions; median 15.5, range 1–32). (b) Distribution of self-reported device heat levels among 176 entries with a recorded setting on the 0–7 scale (median 6, IQR 5–6). (c) Percentage of 188 submitted entries reporting each symptom. One headache and one dry-mouth/skin report were recorded (0.532% each); the other four listed symptoms were not reported. Source values are in Table 1 and Supplementary Tables S15–S16

**Fig. S6** Sensitivity of immune-adjusted predictor effects to the cell reference. (a) Concordance of long-term IAA effect sizes for clock, SystemsAge organ scores, and PhysAge predictors under the primary 12-cell and secondary 19-cell methylation-inferred references (Pearson r=0.6867); purple points identify the eight predictors meeting within-family BH-FDR<0.05 under age/sex-adjusted EAA. (b) EAA, 12-cell IAA, and 19-cell IAA effect sizes for those eight predictors. DNAmCystatinC met 19-cell IAA FDR<0.05 (d=0.82, p=0.000977, FDR=0.00977), whereas none of the eight met FDR<0.05 under the primary 12-cell IAA model. This is a model-sensitivity comparison, not a mediation analysis. Source statistics are in Supplementary Tables S1, S2, and S12

**Fig. S7** Direction-split enrichment of long-term suggestive CpGs. Hyper-and hypomethylated CpGs at raw P<1×10-4 were analyzed separately. (a–b) Probe-bias-corrected gometh KEGG enrichment for hypermethylated and hypomethylated lists, respectively. (c–d) Secondary clusterProfiler GO Biological Process enrichment for hypermethylated and hypomethylated lists, respectively. Point position represents -log10(nominal p), point size represents the number of represented genes, and color represents nominal-p tier; stars identify FDR<0.05. ABC transporters was the direction-split KEGG term meeting FDR<0.05 and arose from the hypermethylated list

**Fig. S8** Sample-collection timeline and physical BeadChip allocation. Forty samples are plotted by day relative to the first study draw, BeadChip barcode, and timepoint. Among the 11 paired long-term participants, TP1 and TP3 samples occupied disjoint chip sets and no participant had both samples on the same chip. The baseline point on a TP3-used chip belonged to an unpaired baseline-only participant and did not provide a paired cross-timepoint comparison. Source metadata are in Supplementary Tables S17a–c

**Fig. S9** Participant-level long-term trajectories for the eight EAA findings meeting within-family BH-FDR<0.05. Age/sex-adjusted residuals are shown at TP1 and TP3 for PCDNAmTL, DepressionBarbu, SystemsAge Blood, SystemsAge Lung, DNAmPulsePressure, DNAmCystatinC, DNAmHDL, and DNAmDHEAS among 11 paired participants. Gray lines connect individual participants; black points and lines show group means. Source trajectories and leave-one-participant-out results are in Supplementary Tables S18a–c

**Fig. S10** Global sensitivity test of methylation-inferred leukocyte composition. Bars show standardized mean TP3-minus-TP1 differences for the 11 leukocyte fractions entered as covariates in the IAA model. The exact paired sign-flip test across all 2,048 sign patterns yielded T=18.788 and p=0.0967. Treg and memory B cells contributed 30.5% and 21.0% of the global statistic, respectively. Source statistics are in Supplementary Tables S19a–c

## Online Resource 2: Supplementary Table Legends

**Table S1** Complete epigenetic clock and SystemsAge organ clock results. Age/sex-adjusted EAA and primary 12-cell IAA statistics are reported for predictors in each contrast, including Cohen’s d, nominal p values, within-family BH-FDR values, and significance tiers.

**Table S2** Complete DNAm PhysAge results. Age/sex-adjusted EAA and primary 12-cell IAA statistics are reported for 10 physiological-trait methylation proxies in each contrast, including Cohen’s d, nominal p values, within-family BH-FDR values, and significance tiers.

**Table S3** Complete EBP v2 metabolite-proxy results. Age/sex-adjusted EAA and primary 12-cell IAA statistics are reported for 1,378 scores in each contrast, including Cohen’s d, nominal p values, within-family BH-FDR values, and significance tiers.

**Table S4** Complete Marioni protein EpiScore results. Age/sex-adjusted EAA and primary 12-cell IAA statistics are reported for 113 protein EpiScores in each contrast, including Cohen’s d, nominal p values, within-family BH-FDR values, and significance tiers.

**Table S5** Complete suggestive DML sets. All CpGs meeting raw P<1×10-4 are reported for the acute and long-term contrasts, with CpG and gene annotation, genomic position, log-fold change on M values, mean beta-value difference, raw P, BACON-calibrated p and FDR, genomic-control p and FDR, and reporting tier.

**Table S6** Complete DML pathway analyses. The table contains enrichment results (acute and long-term) from primary probe-bias-corrected gometh analyses of KEGG and MSigDB Hallmark collections and secondary clusterProfiler analyses of KEGG and Gene Ontology Biological Process, Molecular Function, and Cellular Component collections, with represented-gene counts, nominal p values, FDR values, and reporting tiers.

**Table S7** Complete HMFDR-positive DMR sets. All DMRcate regions meeting HMFDR<0.05 are reported: acute regions and long-term regions with genomic interval, nearest-gene annotation, CpG count, mean beta-value difference, Fisher combined p value, HMFDR, and reporting tier.

**Table S8** Complete long-term DMR-gene pathway analyses. The table contains results derived from genes annotated to the long-term HMFDR-positive regions (GO Biological Process and KEGG terms), with represented-gene counts, nominal p values, FDR values, and reporting tiers. No term met FDR<0.05. Acute enrichment was not performed because only four acute regions had gene annotations.

**Table S9** Reference-cohort contextual comparisons. Age/sex-adjusted standardized differences, nominal p values, and within-family BH-FDR values are reported for predictors at firefighter baseline versus controls and predictors at long-term follow-up versus controls. Firefighter/reference cohort and processing run were aligned, so these comparisons are descriptive.

**Table S10** Acute methylation-inferred leukocyte changes. Paired TP2-minus-TP1 mean changes, Wilcoxon p values, Cohen’s d, and BH-FDR values are reported for 12 inferred leukocyte proportions among acute pairs; no cell proportion met BH-FDR<0.05.

**Table S11** Long-term methylation-inferred leukocyte changes. Paired TP3-minus-TP1 mean changes, Wilcoxon p values, Cohen’s d, and BH-FDR values are reported for 12 inferred leukocyte proportions among long-term pairs. Treg provided the strongest individual signal; no cell proportion met BH-FDR<0.05.

**Table S12** Long-term 19-cell IAA sensitivity analysis. Age/sex-adjusted EAA and 19-cell IAA statistics are reported for clock, SystemsAge organ scores, and PhysAge predictors. Corresponding primary 12-cell IAA results are in Supplementary Tables S1–S2.

**Table S13** EWAS calibration summary. Genomic-control lambda, BACON empirical-null inflation and bias, raw P<1×10-4 counts, raw-BH, genomic-control-FDR and BACON-FDR counts, BACON-calibrated p<1×10-4 counts, and overlap with the raw-P set are reported for the primary long-term no-PC model, long-term with-PC sensitivity model, and acute with-PC model. BACON calibration retained 329 long-term and 126 acute CpGs at p<1×10-4.

**Table S14** Baseline WHO quality-of-life item summary. Mean, standard deviation, and available respondent count are reported for 25 WHO-QoL items scored on a 1–5 Likert scale; item-specific n was 15 or 16.

**Table S15** Participant-linked adherence. Distinct logged session counts are reported for long-term participants using de-identified study IDs.

**Table S16** Reported symptom frequency. The percentage of submitted session entries reporting each of six prespecified symptoms is shown.

**Table S 17a–c** Technical allocation and sample metadata. Table S17a cross-tabulates TP1, TP2, and TP3 sample counts across 10 physical BeadChips. Table S17b reports the TP1 and TP3 chip assignments for all long-term pairs and whether each pair shared a chip (none did). Table S17c provides de-identified sample-level timepoint, chip, array position, relative collection day, and BeadChip identifier for the samples.

**Table S18a–c** Leave-one-participant-out sensitivity and source trajectories for the eight long-term EAA findings meeting within-family BH-FDR<0.05. Table S18a summarizes the full-sample and leave-one-out p-value and effect-size ranges for each predictor. Table S18b contains all 88 refits (11 exclusions for each of 8 predictors). Table S18c contains the corresponding 88 participant-level TP1 and TP3 EAA values used in Figure S9.

**Table S19a–c** Global sensitivity analysis of the 11 methylation-inferred leukocyte fractions entered in the IAA model. Table S19a reports the exact paired sign-flip test. Table S19b reports each cell fraction’s standardized mean difference and contribution to the global statistic. Table S19c reports univariate Wilcoxon p values, Cohen’s d, and BH-FDR cross-checks for the same 11 fractions.

