## Supplementary Figures S1-S10 for "Whole-blood DNA-methylation patterns during portable-sauna use in firefighters: an exploratory, single-arm pilot study"

**Journal:** GeroScience

**Authors:** Varun B. Dwaraka; Adiv A. Johnson; Kirsten Seale; Nolan Kahal; Dua Sheikh; Sean Morrissey; Ryan Smith

### Fig. S1

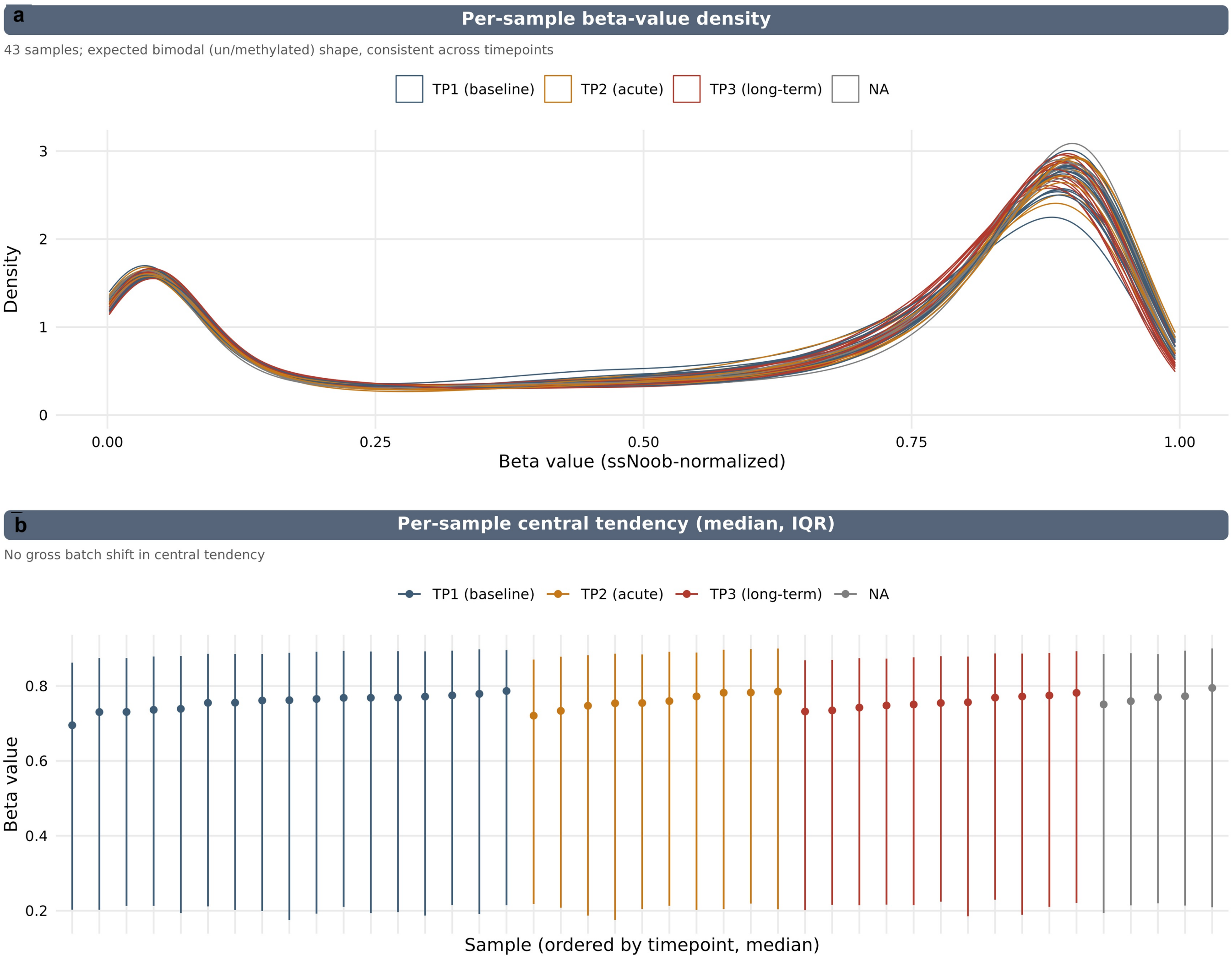


**Fig. S1** Post-normalization DNA-methylation quality control. (a) Per-sample ssNoob-normalized beta-value density distributions, colored by timepoint where available. (b) Per-sample medians and interquartile ranges, ordered by timepoint and median. The expected bimodal methylation distributions and similar central tendencies were retained across samples

**Fig. S2**

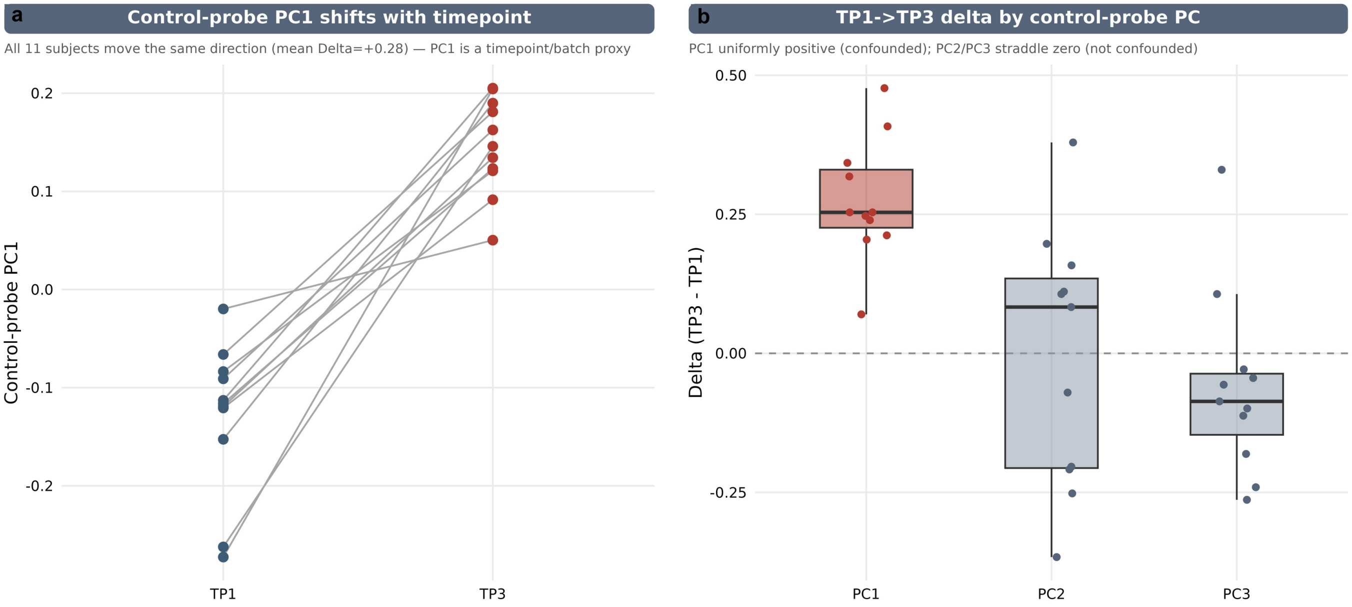


**Fig. S2** Alignment of control-probe PC1 with long-term timepoint. (a) Paired control-probe PC1 values for the 11 TP1–TP3 pairs; every participant shifted in the positive direction (mean TP3-minus-TP1 change, +0.28). (b) Paired changes for control-probe PCs 1–3. PC1 was uniformly positive, whereas PC2 and PC3 included changes on both sides of zero. This alignment motivated use of the no-PC long-term EWAS as the primary descriptive model and the with-PC model as a sensitivity analysis

### Fig. S3

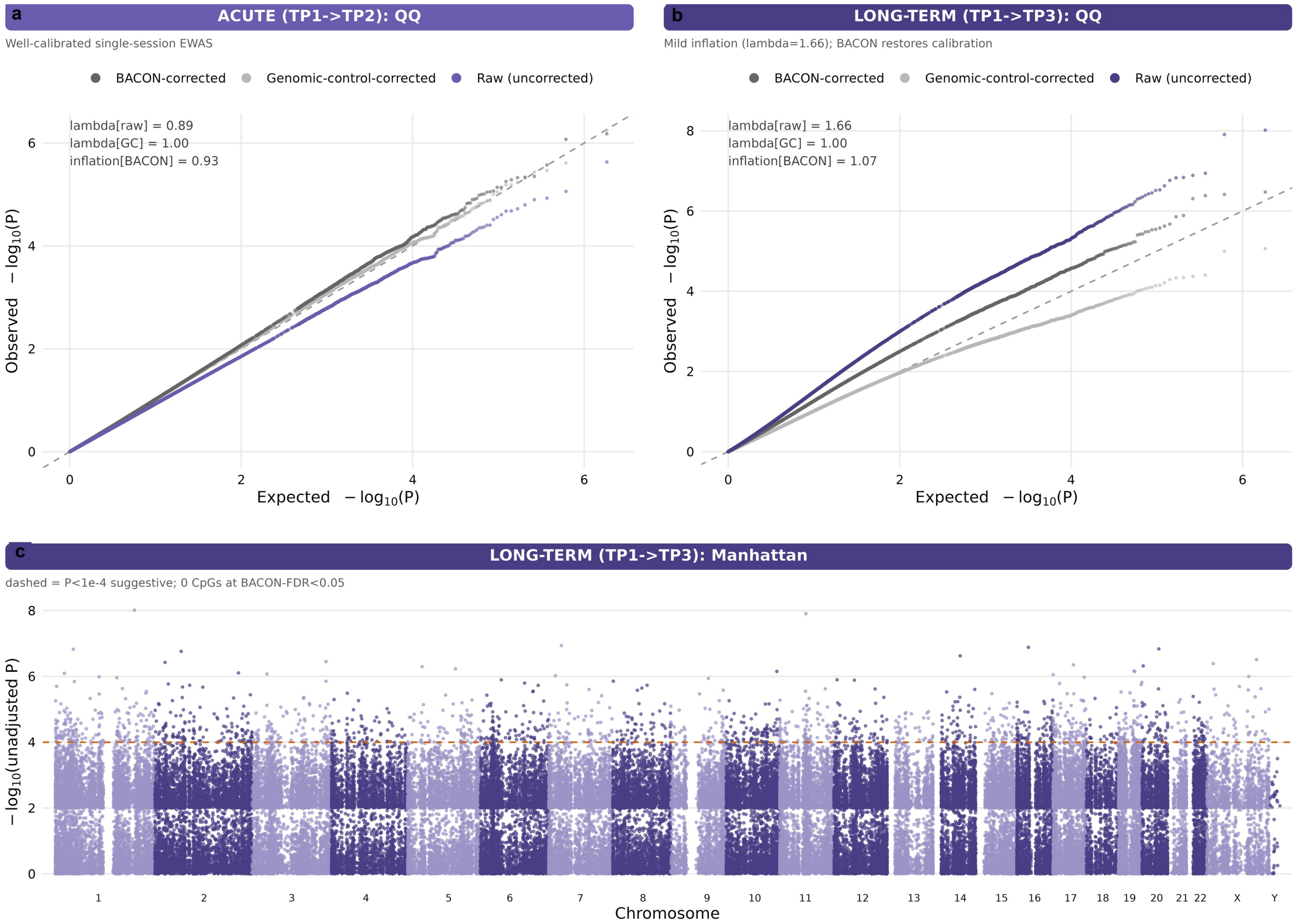


**Fig. S3** EWAS calibration and genome-wide view. Quantile–quantile plots show raw, genomic-control-corrected, and BACON-calibrated p values for (a) the acute TP1–TP2 model (raw lambda=0.890; BACON inflation=0.926) and (b) the primary long-term TP1–TP3 model (raw lambda=1.664; BACON inflation=1.072). (c) Manhattan plot of raw long-term p values across the autosomes and sex chromosomes; the dashed line marks raw P=1×10-4. BACON calibration retained 126 acute and 329 long-term CpGs at p<1×10-4; no CpG met BACON-FDR<0.05. Calibration counts are in Supplementary Table S13

### Fig. S4

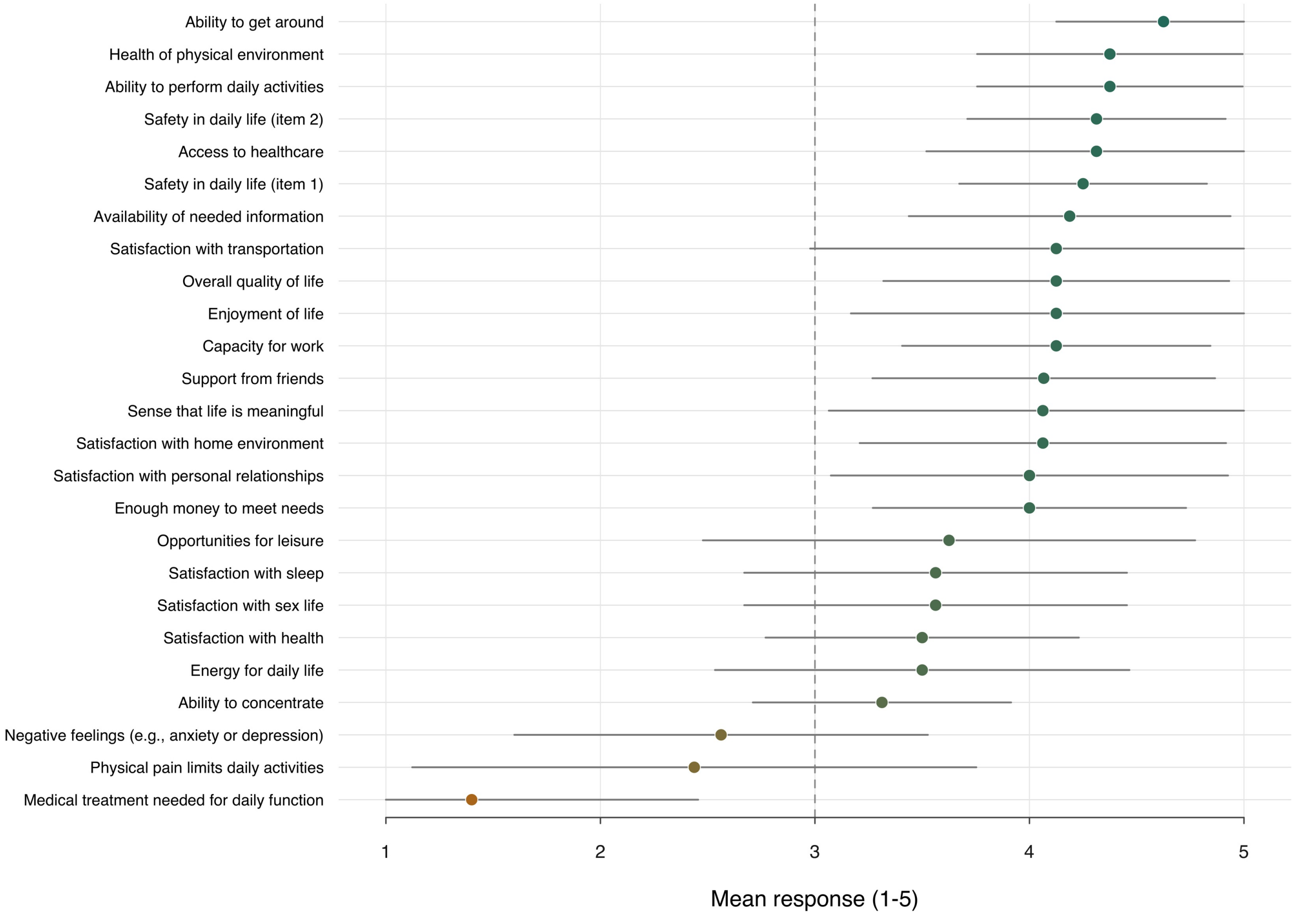


**Fig. S4** Baseline WHO quality-of-life item ratings. Means and standard deviations are shown for WHO-QoL items scored on a 1–5 Likert scale; the dashed vertical line marks the midpoint of 3. Item-specific sample sizes were 15 or 16 respondents. These cross-sectional questionnaire data were not linked to the methylation analyses. Source values are in Supplementary Table S14

### Fig. S5

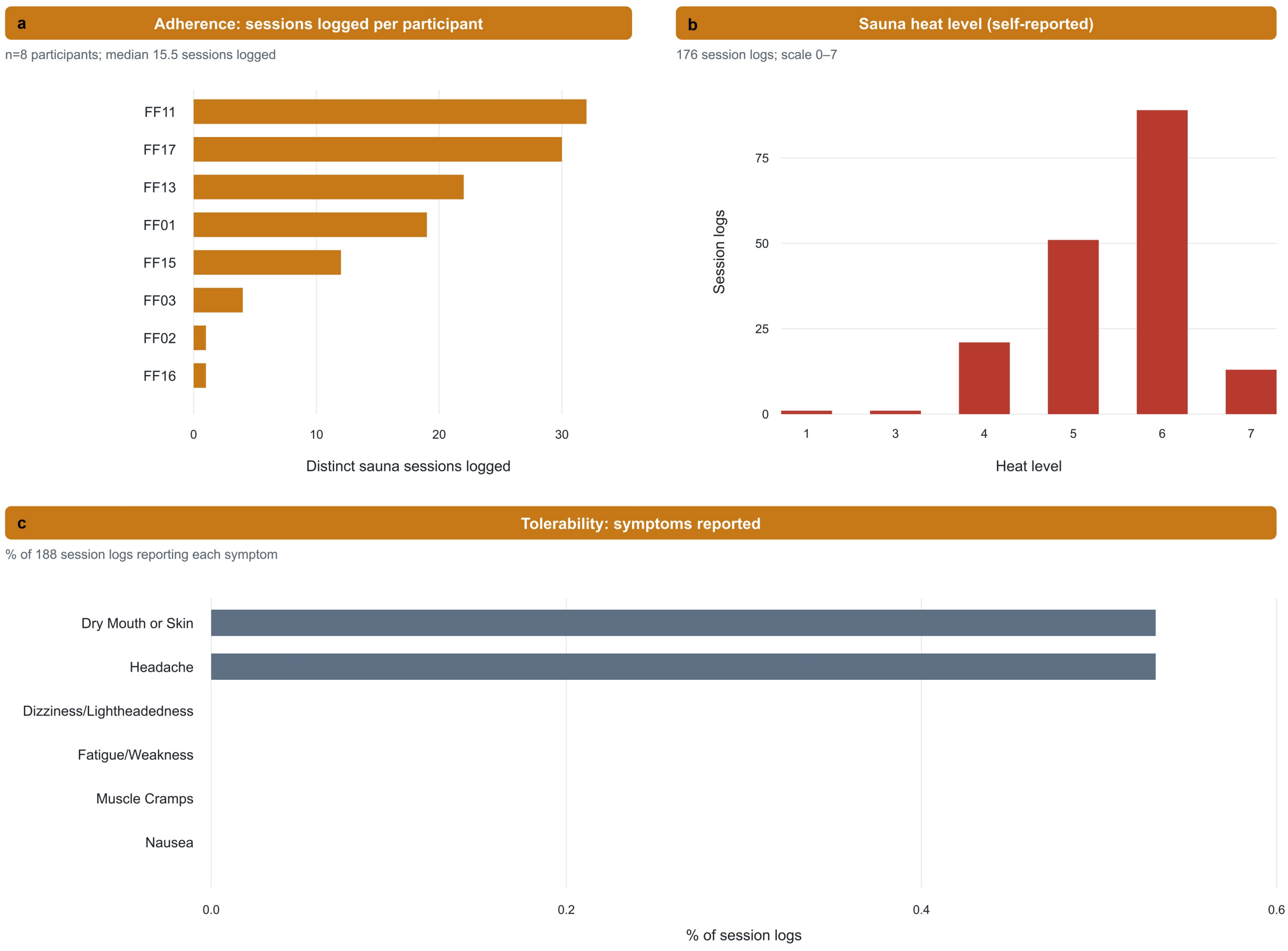


**Fig. S5** Adherence and tolerability from submitted session logs. (a) Participant-linked session counts for 8 of the 11 long-term participants (121 linked sessions; median 15.5, range 1–32). (b) Distribution of self-reported device heat levels among 176 entries with a recorded setting on the 0–7 scale (median 6, IQR 5–6). (c) Percentage of 188 submitted entries reporting each symptom. One headache and one dry-mouth/skin report were recorded (0.532% each); the other four listed symptoms were not reported. Source values are in Table 1 and Supplementary Tables S15–S16

### Fig. S6

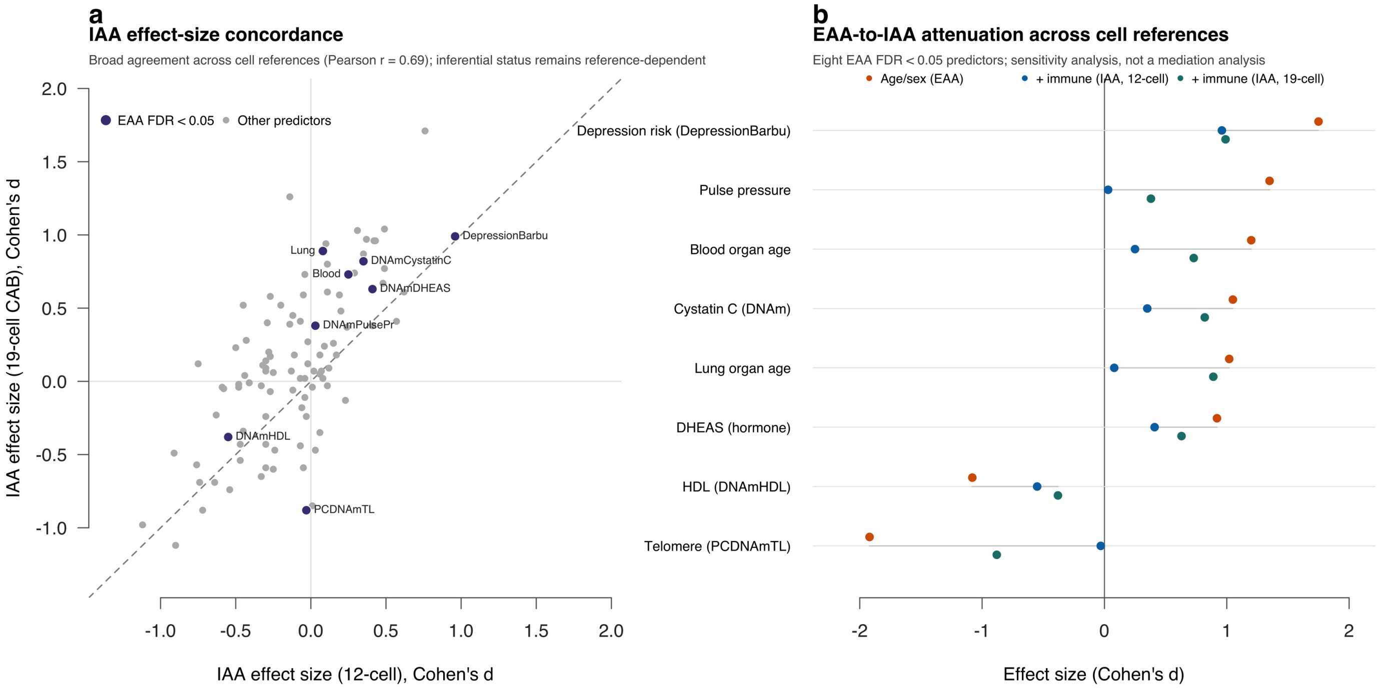


**Fig. S6** Sensitivity of immune-adjusted predictor effects to the cell reference. (a) Concordance of long-term IAA effect sizes for clock, SystemsAge organ scores, and PhysAge predictors under the primary 12-cell and secondary 19-cell methylation-inferred references (Pearson r=0.6867); purple points identify the eight predictors meeting within-family BH-FDR<0.05 under age/sex-adjusted EAA. (b) EAA, 12-cell IAA, and 19-cell IAA effect sizes for those eight predictors. DNAmCystatinC met 19-cell IAA FDR<0.05 (d=0.82, p=0.000977, FDR=0.00977), whereas none of the eight met FDR<0.05 under the primary 12-cell IAA model. This is a model-sensitivity comparison, not a mediation analysis. Source statistics are in Supplementary Tables S1, S2, and S12

### Fig. S7

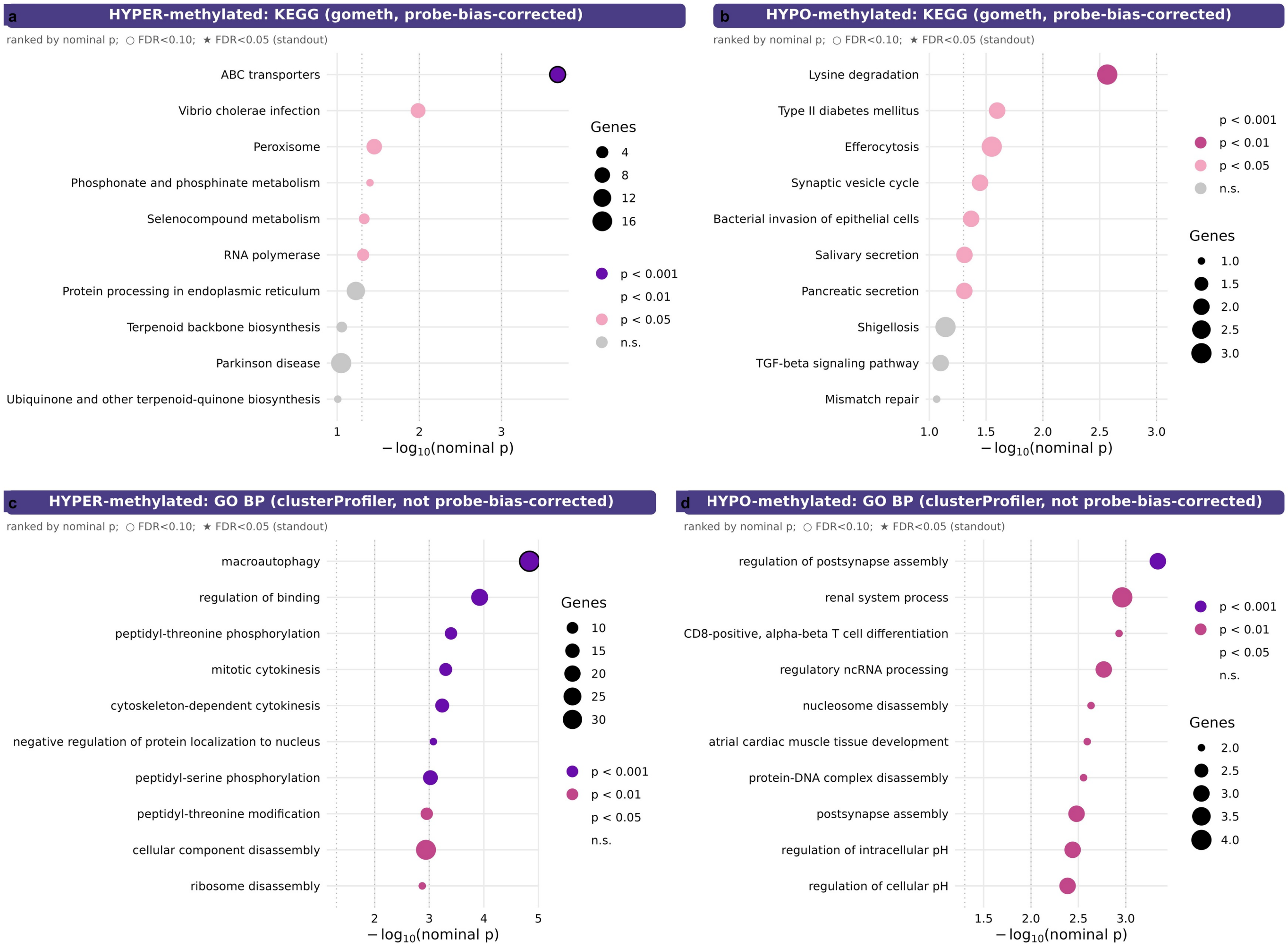


**Fig. S7** Direction-split enrichment of long-term suggestive CpGs. Hyper- and hypomethylated CpGs at raw P<1×10-4 were analyzed separately. (a–b) Probe-bias-corrected gometh KEGG enrichment for hypermethylated and hypomethylated lists, respectively. (c–d) Secondary clusterProfiler GO Biological Process enrichment for hypermethylated and hypomethylated lists, respectively. Point position represents -log10(nominal p), point size represents the number of represented genes, and color represents nominal-p tier; stars identify FDR<0.05. ABC transporters was the direction-split KEGG term meeting FDR<0.05 and arose from the hypermethylated list

**Fig. S8**

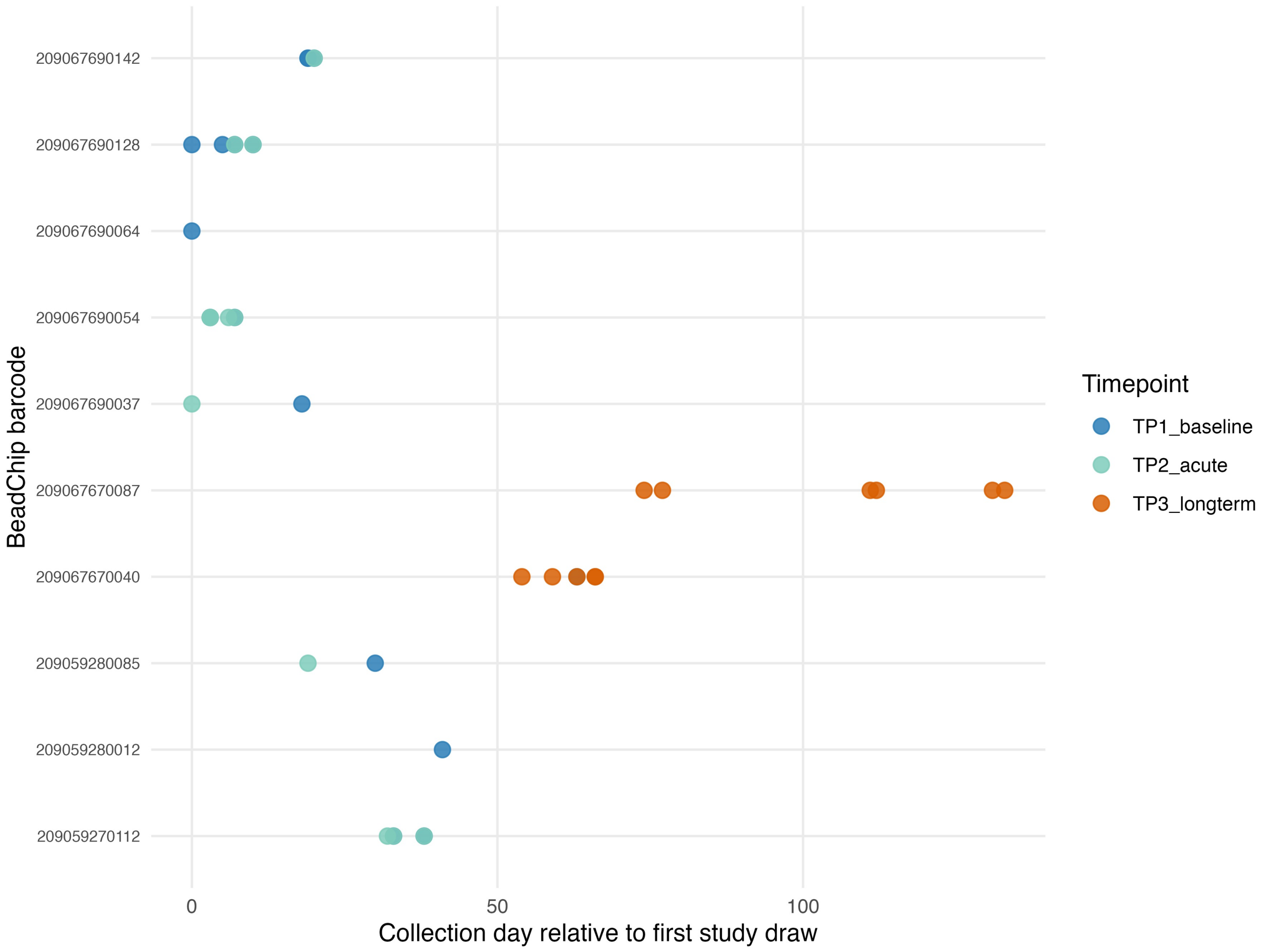


**Fig. S8** Sample-collection timeline and physical BeadChip allocation. Forty samples are plotted by day relative to the first study draw, BeadChip barcode, and timepoint. Among the 11 paired long-term participants, TP1 and TP3 samples occupied disjoint chip sets and no participant had both samples on the same chip. The baseline point on a TP3-used chip belonged to an unpaired baseline-only participant and did not provide a paired cross-timepoint comparison. Source metadata are in Supplementary Tables S17a–c

**Fig. S9**

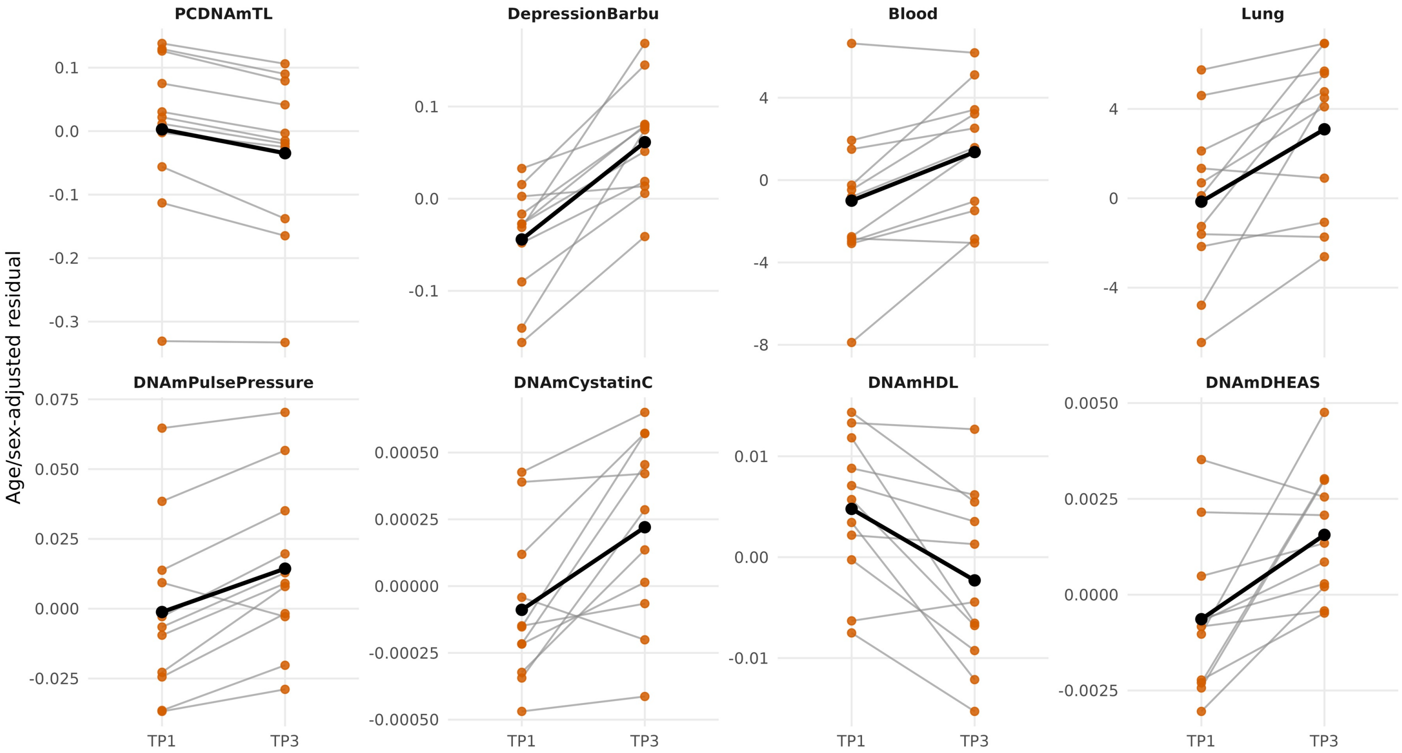


**Fig. S9** Participant-level long-term trajectories for the eight EAA findings meeting within-family BH-FDR<0.05. Age/sex-adjusted residuals are shown at TP1 and TP3 for PCDNAmTL, DepressionBarbu, SystemsAge Blood, SystemsAge Lung, DNAmPulsePressure, DNAmCystatinC, DNAmHDL, and DNAmDHEAS among 11 paired participants. Gray lines connect individual participants; black points and lines show group means. Source trajectories and leave-one-participant-out results are in Supplementary Tables S18a–c

**Fig. S10**

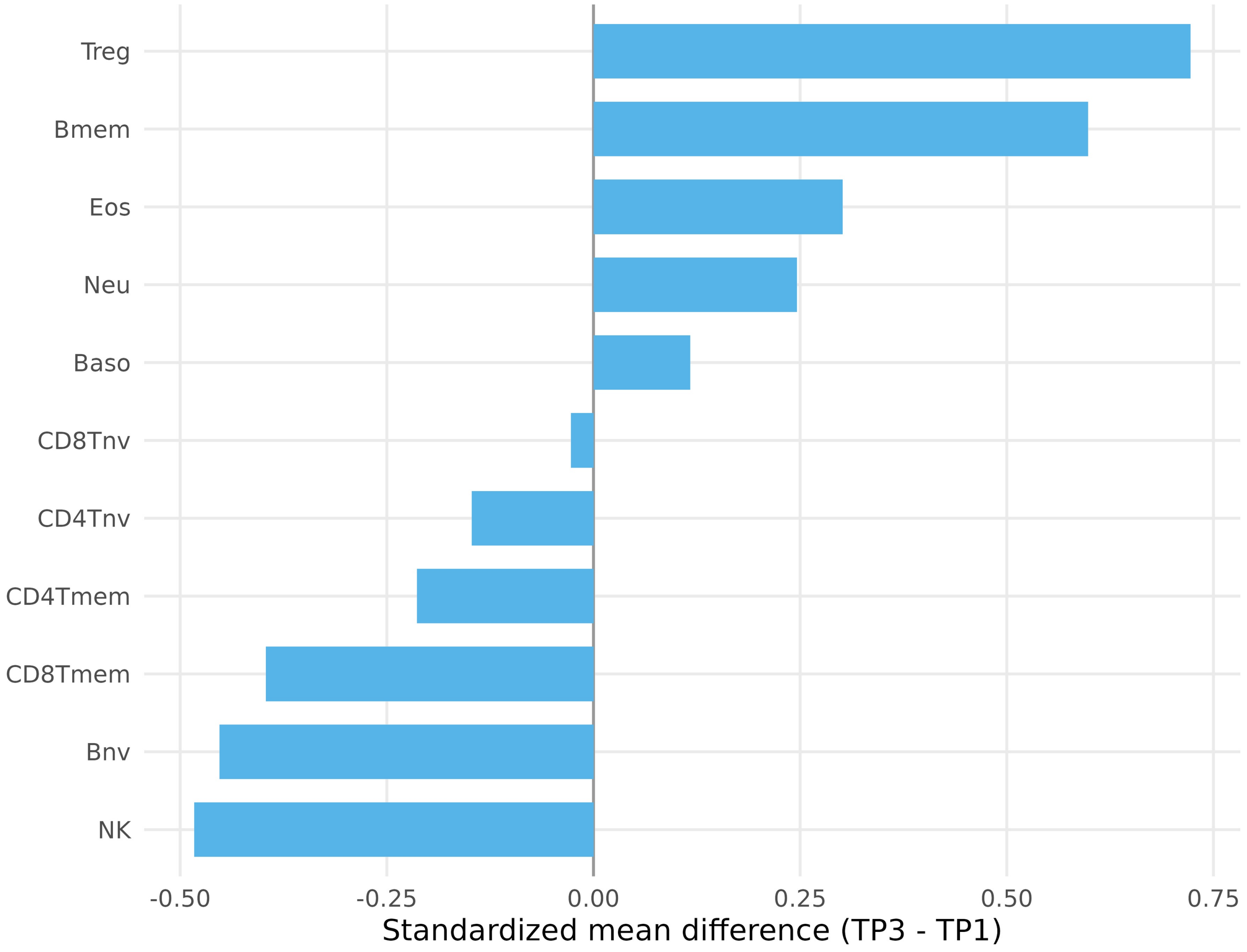


**Fig. S10** Global sensitivity test of methylation-inferred leukocyte composition. Bars show standardized mean TP3-minus-TP1 differences for the 11 leukocyte fractions entered as covariates in the IAA model. The exact paired sign-flip test across all 2,048 sign patterns yielded T=18.788 and p=0.0967. Treg and memory B cells contributed 30.5% and 21.0% of the global statistic, respectively. Source statistics are in Supplementary Tables S19a–c
